# DNA-Directed Assembly of Multivalent Lipid Nanoparticles for Targeted T Cell Gene Delivery

**DOI:** 10.1101/2025.09.04.674323

**Authors:** Mary D. Kelly, Timothy Q. Vu, Atiriya U. Iyer, Yiming Luo, Aiden P. Linderman, Lariana Cline, Crystal Sanchez, Neha P. Kamat

**Affiliations:** Center for Synthetic Biology, Northwestern University, Evanston, IL, 60208

## Abstract

Lipid nanoparticles (LNPs) are a powerful emerging tool for *in vivo* T cell engineering with applications ranging from B cell lymphomas to other cancers and autoimmune diseases. Key challenges in designing these therapeutics include achieving both precise cell targeting and efficient mRNA translation. While single-targeted LNPs have been extensively studied, bispecific LNPs have only been briefly explored. Engagement of multiple T cell receptors offers the opportunity for enhanced mRNA delivery, expression, and T cell targeting. Here, a DNA-tethering method enables rapid modification of lipid nanoparticles with commercial antibodies. Using this strategy, we evaluated a variety of bispecific LNPs for targeted mRNA delivery to T cells both *in vitro* and *in vivo*. We identify bispecific formulations that improve targeting and subsequent transfection of T cells *in vitro* and *in vivo* relative to monotargeted LNPs. Additionally, we find that targeting molecules can alter LNP biodistribution to the spleen and liver. This fast and efficient approach to assembling antibody-targeted LNPs should enable high-throughput screening of diverse antibody combinations for improved specificity and efficiency of *in vivo* gene delivery.

**Significance Statement:** A major challenge in *in vivo* gene delivery is achieving both precise targeting and efficient mRNA translation. Multitargeted LNPs offer a potential solution to this challenge; however, their rapid assembly remains difficult, necessitating the development of new methods to construct and evaluate targeted LNPs. Here, we use a DNA-tethering method to enable rapid antibody modification of LNPs and evaluate bispecific formulations for targeted T cell mRNA delivery *in vitro* and *in vivo*. We find bispecific LNPs improve T cell targeting and expression compared to single-targeted particles. To the best of our knowledge, this study represents the first systematic screening and comparison of bispecific LNPs. Our method provides a modular approach for identifying effective antibody combinations to enhance *in vivo* gene delivery that can be customized for different undruggable diseases.

## Introduction

Lipid nanoparticles (LNPs) have emerged as a potent and tunable platform for T cell engineering with vast potential to transform treatments of cancers, T cell infections, and autoimmune diseases (1). For the treatment of B-cell lymphomas, chimeric antigen receptor (CAR) T therapies have been highly effective through laborious and expensive process of engineering T cells *ex vivo* (2). There has been a shift towards *in vivo* generation of CAR T cells, which could be performed with off-the-shelf therapeutics that will streamline CAR T cell production and increase treatment accessibility. *In vivo* CAR**’**s have reached clinical trials for treating B-cell lymphomas, as well as autoimmune diseases, multiple myeloma, epithelial tumors, and liver cancer (3-5).

Key challenges for *in vivo* T cell gene therapies lie in the development of delivery vehicles capable of both highly efficient and on target T cell transfection (2). LNP delivery approaches are currently limited to localized administration and hepatic cells, which have a natural propensity for particle uptake. To overcome this limitation, two main strategies have been pursued to redirect LNP biodistribution: (1) optimizing the structure of ionizable lipids to achieve tropism for non-hepatic cells and organs, and (2) surface-modifying LNPs with receptor-specific antibodies for targeted delivery (1, 6, 7). The latter approach has been explored using various single T cell receptor targets, including CD3 (8, 9) CD4 (10), CD5 (8, 11), CD7 (8), and CD8 (12). These formulations generally result in low transfection efficiency which may be attributed to poor or partial T cell activation, which has been shown to be key for the expression of LNP-delivered mRNA (13). In the body, T cell activation is controlled by interactions with antigen presenting cells (APCs). Activation is a multivalent phenomenon, achieved through providing a primary activation signal, usually through engagement with the T cell receptor complex (TCR) in combination with a costimulatory signal (14). Most often, a combination of antibodies against CD3 and CD28 are used to activate T cells *ex vivo* to mimic the stimulation provided by APCs (15). In previous work, bispecific LNPs modified with a combination αCD3 and αCD28 antibodies have been explored for *ex vivo* T cell engineering and showed an increase transfection in comparison to LNPs modified with αCD3 alone (13). Some bispecific formulations such as αCD3/αCD7-LNPs have also seen success for *in vivo* CAR generation to treat B cell leukemia (16). Further screening of bispecific LNPs both *ex vivo* and *in vitro* could identify new T cell surface targets that increase efficacy of mRNA delivery. An understanding of how co-targeting use of immunomodulatory signals impacts T cell transfection is key for therapeutic optimization and requires methods to rapidly assemble and screen many combinations of targeted LNPs (tLNPs).

Several technical barriers in antibody conjugation have limited progress in development of tLNPs. Traditional methods for conjugating antibodies to the surface of LNPs such as amide formation and maleimide chemistry are often time-consuming, and result in low efficiency, and lead to incorrect antibody orientation which is not conducive for screening large combinations of targeting molecules (17). While click chemistry offers a rapid method to attach multiple antibodies, it lacks sufficient orthogonal attachment chemistry to control the ratios or spatial arrangement of attached proteins (18, 19). In contrast, oligonucleotide assembly has been shown to be a robust, rapid, and precise method for tethering of antibodies to nanoparticles (20). By leveraging the highly specific base-pairing interactions between complementary nucleic acid sequences, researchers can design and control specific material and biomolecule interactions (21-25).

Here, we developed a workflow for the rapid modification of LNPs using DNA-tethering of commercially available antibodies for screening combinations of antibodies for enhanced transfection efficiency in T cells. T cells were chosen as a model due to their therapeutic relevance, resistance to transfection, and the well-characterized expression of specific surface markers across their subsets. We demonstrate the use of bispecific LNPs for both *in vitro* delivery to Jurkat and primary T cells and *in vivo* screening of top-performing formulations. By leveraging this DNA-anchoring method for antibody conjugation to LNPs, we can rapidly identify antibody combinations capable of efficiently targeting and transfecting T cells, paving the way for the development of highly effective *in vivo* T cell engineering for multiple pathologies.

## Results

### DNA-tethering enables rapid and efficient attachment to LNP surface

A DNA-tethering conjugation method was used to attach antibodies to the surface of LNPs16, 20 (Fig. 1*A*). First, LNPs were synthesized using the ionizable cationic lipid Dlin-MC3-DMA, which was also used in the first FDA-approved LNP gene therapy (26). Oligonucleotide-conjugated lipid (DNA-lipid) was synthesized using copper-free click chemistry (25) and confirmed using mass spectrometry (*SI Appendix*, Fig. S1). A complementary strand of ssDNA was attached to the antibodies of interest (DNA-Ab) using a photoactivated Protein G which enables site specific modification of nearly all IgG antibodies with high efficiency (AlphaThera) (27). Successful DNA-Ab synthesis was confirmed using gel electrophoresis (*SI Appendix*, Fig. S2). The DNA-lipid was post-inserted into preformed LNPs and the insertion was assessed using a fluorescent complementary DNA (cDNA) followed by size exclusion chromatography (SEC) (Fig. 1*B-D* and *SI Appendix*, Fig. S3). We compared the use of a DNA-lipid strand containing a 15 atom tetraethylene glycol (TEG) spacer to a DNA-lipid without the spacer (*SI Appendix*, Table S1). We found a greater than 2-fold increase in the average amount of conjugated cDNA using the DNA-lipid without the TEG spacer (Fig. 1*B*). This difference in conjugation efficiency indicates that the TEG spacer either hinders the lipid post-insertion process or sterically blocks the binding of the cDNA strand. All further studies were completed without the TEG spacer.

**Figure 1.**
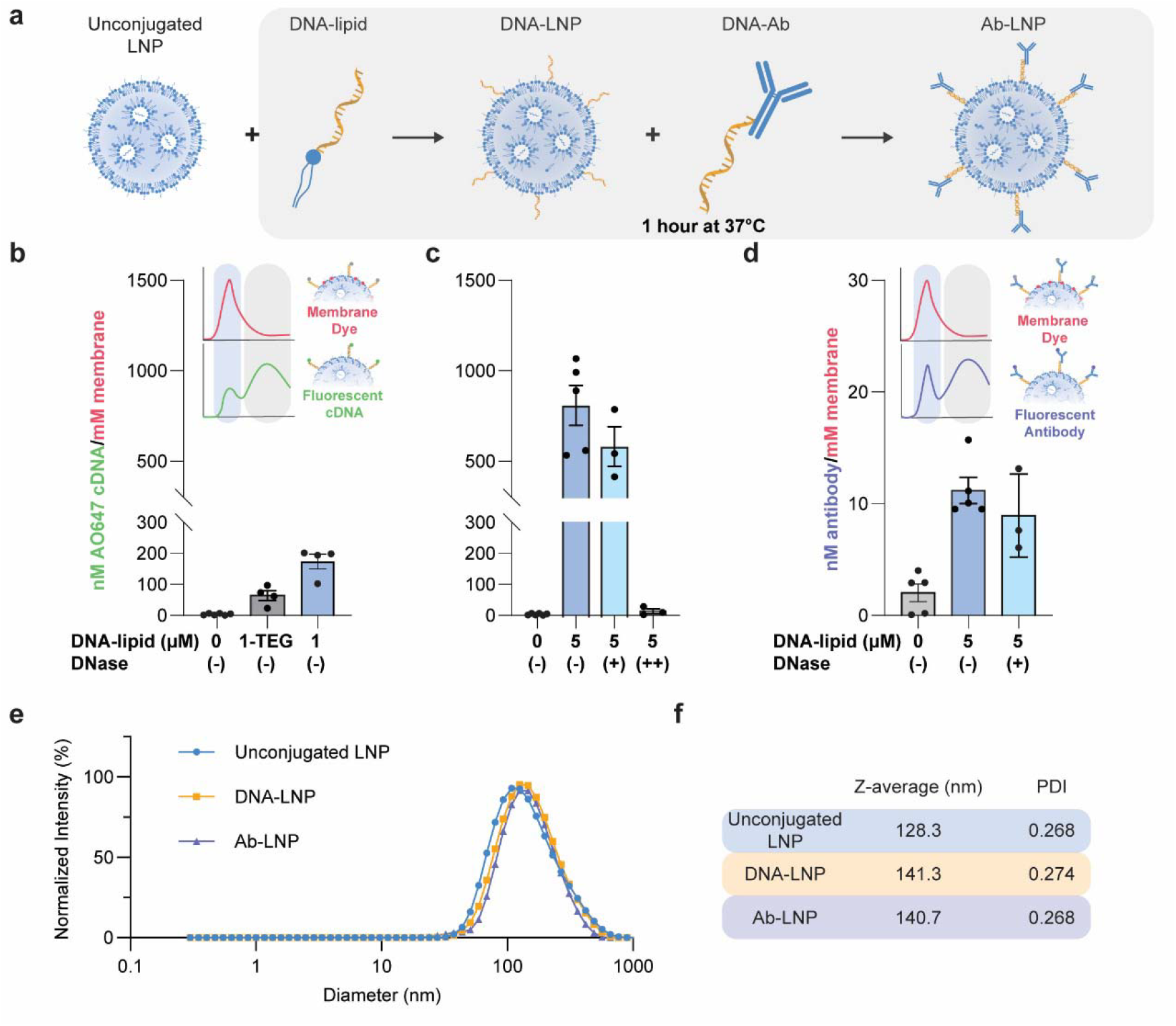
Method for DNA-tethering of antibodies to LNPs and particle characterization. (a) Schematic of modification of LNPs through a 1-hour incubation of LNP, DNA-lipid, and DNA-Ab at 37°C. (b) DNA-LNPs were incubated with a fluorescent Atto647 cDNA strand and run on SEC. The nM cDNA per mM of LNP membrane were quantified. DNA-lipid was added at a concentration of 1 uM with and without a TEG spacer (error bars = standard error of mean). (c) DNA-lipid was added at a concentration of 5 uM. The modified particles were incubated for 30 min at 37°C with 0.5 U/mL (+) and 10,000 U/mL (++) DNase I before SEC (error bars = standard error of mean). (d) DNA-LNPs were incubated with a fluorescent AF647 DNA-Ab and run on SEC. The nM of antibody per mM of LNP membrane was quantified. This was also measured after incubation with 0.5 U/mL DNase I (error bars = standard error of mean). (e) Particle size was measured using DLS for LNPs before and after modification (n = 2). (f) The z-average and PDI for the unconjugated LNPs, DNA-LNPs and tLNPs are shown (n = 2).

We next wanted to assess the stability of our DNA-tethers in the presence of physiologically relevant deoxyribonuclease (DNase) concentrations. For this modification method to be useful *in vivo*, it would be necessary for the attachment strategy to remain stable. A DNase I concentration of 0.5 U/mL was used, which falls slightly above the average DNase activity in human plasma reported by Tamkovich et al. (28) of 0.356 ± 0.410 U/mL. As a positive control for DNase I activity a concentration of 10,000 U/mL was used to confirm DNase activity. It was found that at the concentration of 0.5 U/mL, the cDNA strands remained attached to the particle surface whereas at high DNase concentrations of 10,000 U/mL the cDNA was almost completely cleaved as assessed by SEC (Fig. 1*C*).

Following this, we quantified the attachment of the DNA-Ab to the surface of our particle. We used 5 µM of DNA-lipid and 0.06 µM of fluorescent AF647 CD28 DNA-Ab to modify our LNPs. After 30 min, these synthesized particles were run on SEC and the average nM of antibody per mM of LNP lipid was found to be 10.7 ± 1.3 (Fig. 1*D*). This value did not change significantly after a 30 min incubation period with 0.5 U/mL DNase I, indicating that these particles have the potential to be stable *in vivo* (20). Using NanoSight300 (Malvern), it was determined across 4 LNP batches containing either mCherry or firefly luciferase (Fluc) mRNA that the concentration was 6.33×10^11^ ± 1.2 ×10^11^ particles/mL (*SI Appendix*, Fig. S4 and Table S2). Using the concentration of antibody from SEC in combination with this concentration of particles, it can be estimated that there are approximately 33 ± 3 antibodies per LNP which is approximately 58% of the theoretical maximum amount of antibody added to solution.

The size and encapsulation efficiency of the synthesized LNPs were also characterized to ensure LNP formation. Using DLS the particle size of mCherry LNPs was estimated to be 128.3±1.1 nm with a PDI of 0.27 (Fig. 1*E*). The measured diameter increased to approximately 141.3±2.3 nm and 140.6±1.9 nm for the DNA-lipid modified LNP and the tLNP respectively. The PDI remained consistent (Fig. 1*F*). Further confirmation of approximate LNP size and spherical shape was performed using CryoEM (*SI Appendix*, Fig. S5). The encapsulation efficiency of the mCherry LNPs was 80.2 % using the Quant-it RiboGreen RNA Assay Kit.

### DNA-mediated antibody conjugation to LNPs improves transfection of Jurkat Cells

For initial evaluation of LNP transfection *in vitro*, Jurkat cells were transfected using single-targeted and bispecific LNPs. All Jurkat cell experiments were performed with LNPs encapsulating mRNA encoding for firefly luciferase. We tested the transfection levels and viability of Jurkat cells as a function of mRNA dose for five LNP variations conjugated with αCD3 to establish the functionality of the conjugation strategy (Fig. 2*A*). The potency of αCD3-tLNPs has been well established in literature (8, 9, 13) so this targeting molecule was used as a model in initial validation studies and as a control throughout all subsequent experiments. Overall, Fluc expression increased with LNP dose (Fig. 2*B*). Notably, the addition of DNA to the surface of the particles, the conjugation of an isotype, nonbinding antibody, and an unconjugated LNP with soluble αCD3 in solution all increase transfection relative to an unmodified LNP. The highest luciferase expression, however, occurred when cells were transfected with LNPs with the αCD3-LNP. The lowest viability also occurred in this condition (Fig. 2*C*). To avoid high levels of cell toxicity, a mRNA dose of 750 ng/100,000 cells was used for all subsequent Jurkat cell experiments.

**Figure 2.**
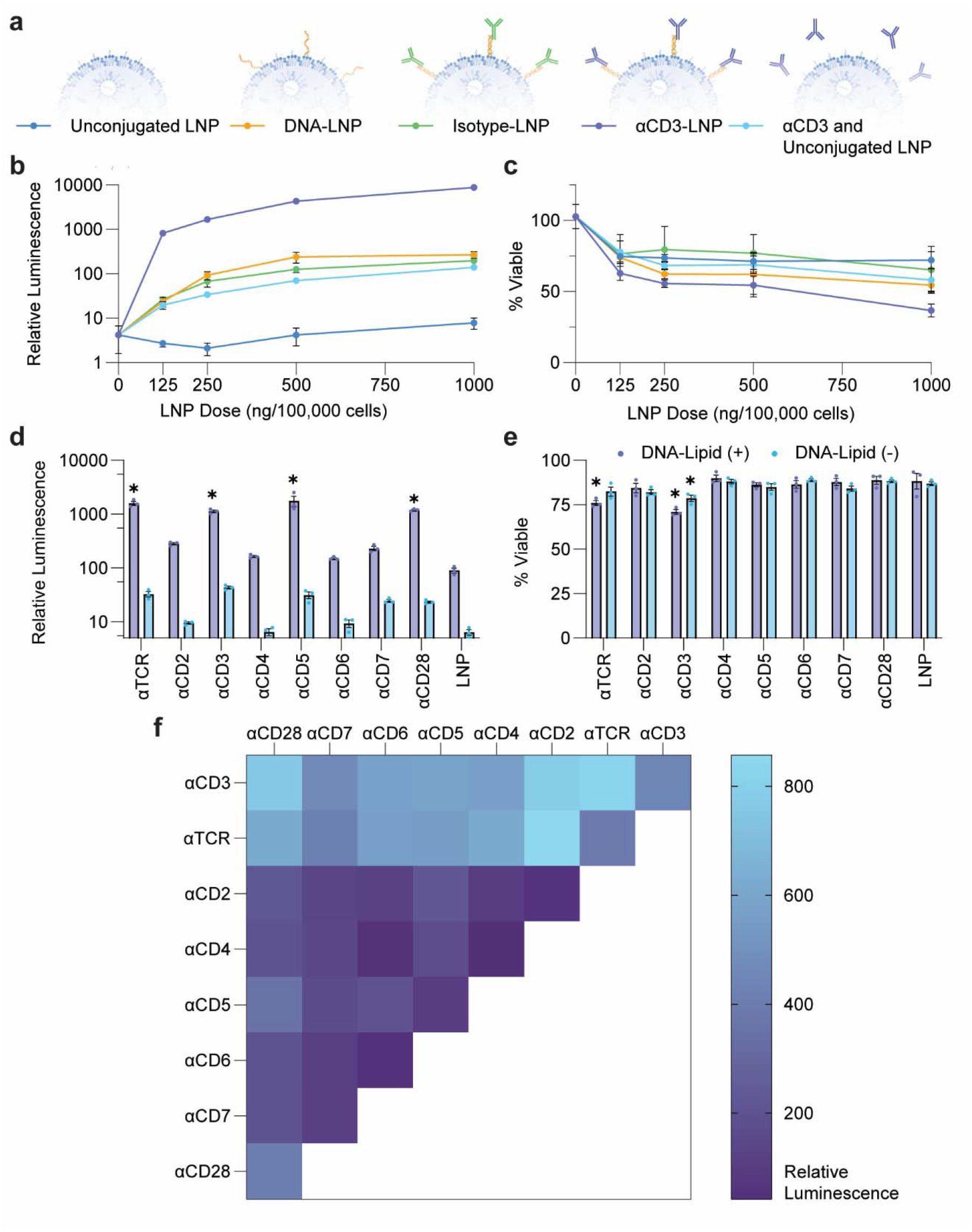
mRNA dose response curve and testing of monospecific and bispecific DNA-tethered tLNPs. (a) Schematic of LNPs tested in the dose response. (b) Luciferase expression of Jurkat cells treated with luciferase-encoding mRNA using a range of mRNA doses after 24 hours (n = 3 replicates, error bars = standard error of the mean). (c) Viability of Jurkat cells treated with luciferase-encoding mRNA using a range of mRNA doses after 24 hours (n = 3 replicates, error bars = standard error of the mean). (d) Luciferase expression in Jurkat cells after treatment with tLNPs with and without the DNA-lipid attachment after 24 hours with a dose of 750 ng/100,000 cells. (e) Viability of Jurkat cells after treatment with tLNPs with and without the DNA-lipid attachment after 24 hours with a dose of 750 ng/100,000 cells. (n = 3 replicates, error bars = standard error of the mean). Statistical analysis included 2-way ANOVA with Dunnett**’**s multiple comparisons test, *p-value < 0.05 as compared to LNP. (f) Heat map of Jurkat luciferase expression after 24 h treatment with bispecific LNPs at a dose of 750 ng/100,000 cells.

Next, antibodies targeting individual common T cell surface receptors were screened for their capacity to increase LNP-mediated transfection of Jurkat cells. Each formation was tested with and without the DNA-lipid to confirm that the conjugation strategy was necessary for high transfection. In each case, the presence of the DNA-lipid alone on an LNP improved Jurkat cell transfection (Fig. 2*D*). The highest T cell transfection was seen with αTCR, αCD3, αCD5 and αCD28-LNPs. Notably, both αTCR and αCD3 engage with proteins which are part of the T cell receptor complex while aCD28 is a commonly used costimulatory molecule. Jurkat cell viability remained stable at approximately 75% for all the antibodies screened with lowest viability seen with αTCR and αCD3-LNPs (Fig. 2*E*).

We next determined if the combinatorial use of these antibodies could improve the transfection of the Jurkat cells. Using a 1:1 ratio of two antibodies we screened 36 different combinations of the antibodies on the surface of LNPs. We found that combinations that included either αTCR or αCD3 seemed to transfect the Jurkat cells with luciferase more efficiently than other combinations. Notably αTCR/αCD2, αCD3/αCD2, αCD3/αTCR, and αCD3/αCD28-LNP had the highest luciferase expression of the 36 combinations screened (Fig. 2*F* and *SI Appendix*, Fig. S6). For future studies, αCD3/αTCR, αCD3/αCD2, and αCD3/αCD28-LNPs were chosen for screening with primary blood cells due to their high transfection in Jurkat cells. In addition, αCD3/αCD4-LNPs were chosen because of the potential to selectively target CD4^+^ T cells which are relevant to CAR T cell and autoimmune cell therapeutics and αCD3/αCD5-LNPs were chosen due to the potential of αCD5 engagement to improve alter T cell function (10, 11).

### Bispecific LNPs enhance transfection of T cells in a human mixed blood cell population through the specific engagement of antibody domains with their respective receptors on T cells

We next evaluated the capacity of bispecific LNPs to specifically transfect T cells in a human mixed blood cell population. LNPs modified with αCD3/αTCR, αCD3/αCD2, αCD3/αCD4, αCD3/αCD5, and αCD3/αCD28 were compared against unmodified LNPs and αCD3-LNPs. Each of these LNP groups were formulated encapsulating mRNA encoding mCherry. Peripheral blood mononuclear cells (PBMCs) were dosed with LNPs at 2000ng mRNA/100,000 cells and incubated for 24 h. Using flow cytometry, we evaluated mCherry expression in T cells (CD3^+^, CD4^+^ helper T cells, and CD8^+^ cytotoxic T cells), CD19^+^ B cells, and CD14^+^ monocytes (*SI Appendix*, Fig. S7). For CD3^+^ cells, which includes subtypes CD4^+^ and CD8^+^, treatment with the monospecific αCD3-LNPs led to transfection of approximately 2.9% of CD4^+^ cells, while the addition of bispecific LNPS containing αCD3 and another antibody (αTCR, αCD2, αCD4, αCD5, and αCD28) led to significantly more transfection except for the αCD3/αCD4-LNPs (Fig. 3*A*). When examining CD4^+^ helper T cells, we observed the bispecific LNP modified with αCD3/αCD4 led to the highest transfection efficiency at ∼8% relative to the nanoparticles which lacked an αCD4 targeting ligand. CD8+ displayed no significant difference in transfection efficiency upon incubation with the different LNP types and an overall lower transfection efficiency than observed with the CD3^+^ and CD4^+^ cell types (Fig. 3*B*). This result suggests that the enhanced transfection efficiency observed in CD3^+^ cells arose from the presence of CD4^+^ targeting molecules and costimulatory binders, which includes both CD2^+^ and CD28^+^.

**Figure 3.**
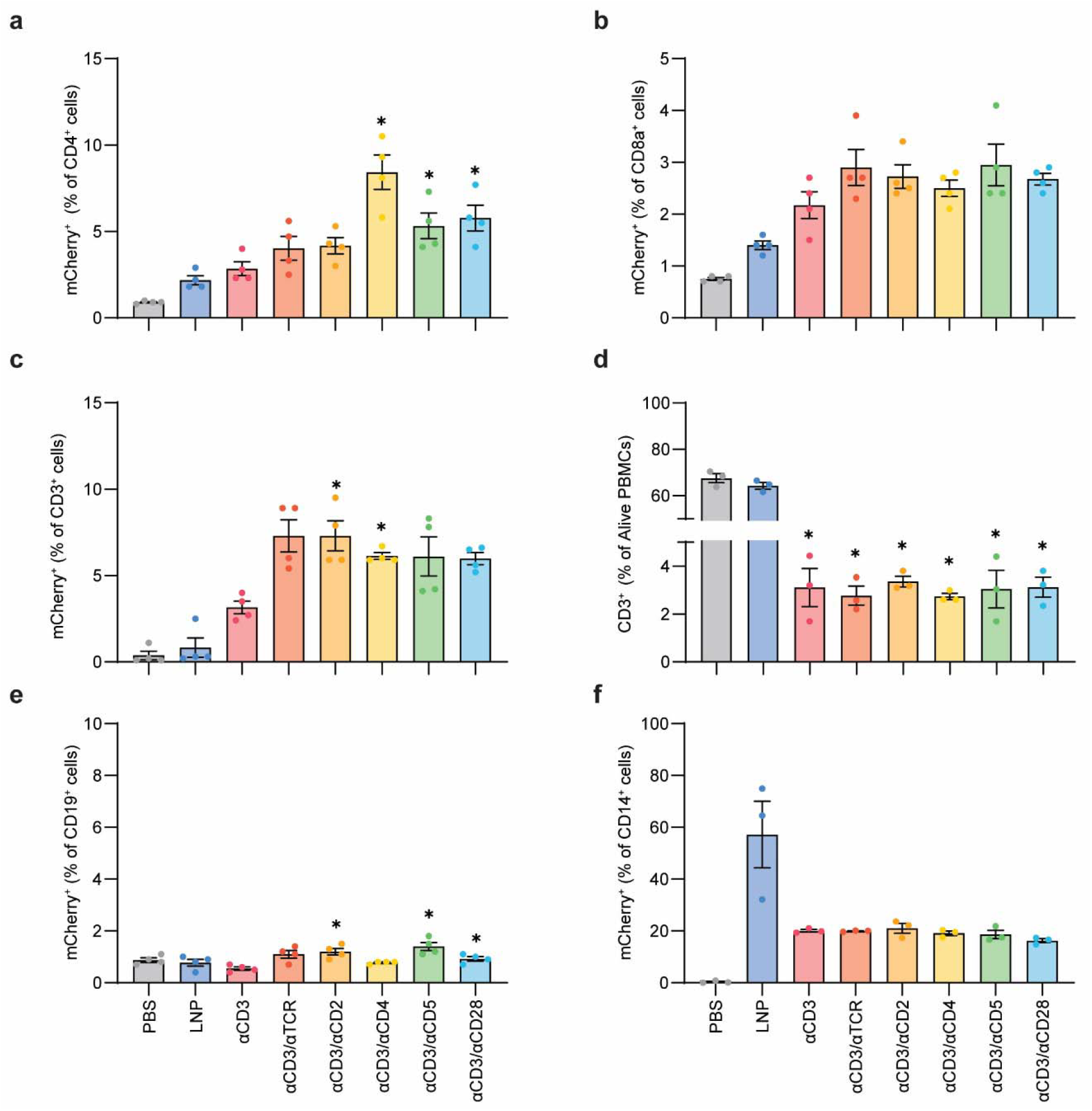
Bispecific LNPs enhance transfection of primary T cells in a human mixed blood cell population *in vitro*. Flow cytometry reveals mCherry+ CD4+ (a), mCherry+ CD8a+, and (b) mCherry+ CD3+ (c) T cells as a percentage of CD4+, CD8+, and CD3+ T cells respectively 24 hours after human PBMCs are treated with PBS or mCherry-mRNA containing unmodified LNP, or tLNPs at a dose of 2000 ng/100,000 cells. (d) CD3+ T cells as a percentage of total alive cells in human PBMCs 24 hours after treatment. mCherry+ CD19+ cells (e) and mCherry+ CD14+ cells as a percentage of CD19+ (b cell marker) and CD14+ (macrophage cell marker) respectively 24 hours after human PBMCs treatment. For (a-c,e) n = 4 biological replicates and for (d,f) n = 3 biological replicates, error bars = standard error of the mean. Statistical analysis included RM one-way ANOVA with multiple comparisons, *p < 0.05 as compared to αCD3-LNPs (a-c, e) and as compared to LNP (d, f).

Consistent with receptor-mediated transfection, we observed downregulation of CD3^+^ receptors when T cells were incubated with LNPs targeted those molecules (Fig. 3*D*). In control groups treated with PBS or unmodified LNPs, approximately 65% of cells were CD3^+^, whereas after treatment with modified LNPs, CD3^+^ expression decreased to approximately 3% of all cells (Fig. 3*D*). The observed downregulation indicates that tLNPs engage in specific antibody-receptor interactions with the intended cell surface receptors, providing evidence that molecularly precise binding underlies the enhanced transfection efficiency

We next evaluated off target uptake by other cell types, which is a concern in the design of targeted delivery vehicles. Both CD19^+^ B cells and CD14^+^ monocytes were evaluated to analyze the potential for off target transfection of the tLNPs. All LNPs resulted in low mCherry expression in CD19^+^ cells, however, the highest mCherrry expression occurred when using αCD3/αCD5-LNPs with ∼1.5% of B cells transfected (Fig. 3*E*). This may be attributed to the dim expression of the CD5 scavenger receptor that has been shown to exist on human B cells and is often used to identify B1 and B2 cell subsets (29). The average monocyte transfection efficiency decreased substantially when using tLNPs compared to untargeted LNPs, dropping from approximately 57% to 20% across all antibody-conjugated LNP groups (Fig. 3*F*). Although tLNPs still exhibited moderate transfection efficiency in monocytes, around 20%, which remains relatively high compared to T cells, this reduction is a promising indication of increased selectivity. Since monocytes and macrophages are major contributors to nonspecific nanoparticle clearance, high uptake by these cells is typically undesirable when the goal is to deliver mRNA to other target cell types, such as T cells. Our findings demonstrate that incorporating T cell-specific targeting ligands onto LNPs not only enhances transfection of the intended cell population but also minimizes nonspecific uptake by non-target immune cells, thereby improving the delivery specificity of mRNA-loaded nanoparticles.

### Antibody conjugated LNPs facilitate organ specific transfection in the spleen

Finally, we evaluated our bispecific LNPs *in vivo*. For this investigation, we used tLNPS that resulted in the highest transfection of T cells *in vitro*: LNPs conjugated with αCD3/TCR, αCD3/αCD2, and αCD3/αCD4-LNPs and compared them to control groups including PBS, unmodified LNPs, an isotype control (iso-LNPs), and single antibody αCD3-LNPs. All LNPs contained a mRNA encoding Fluc and each of the LNP groups were administered to mice at a dose of 0.6 mg/kg of body weight. At time points of 0.5 h, 5 h, and 24 h images were taken using an *in vivo* imaging system (IVIS) to capture luminescent expression over time indicating functional delivery of mRNA and at 24 h organs were collected and imaged individually (Fig. 4*A* and *SI Appendix*, Fig. S8).

**Figure 4.**
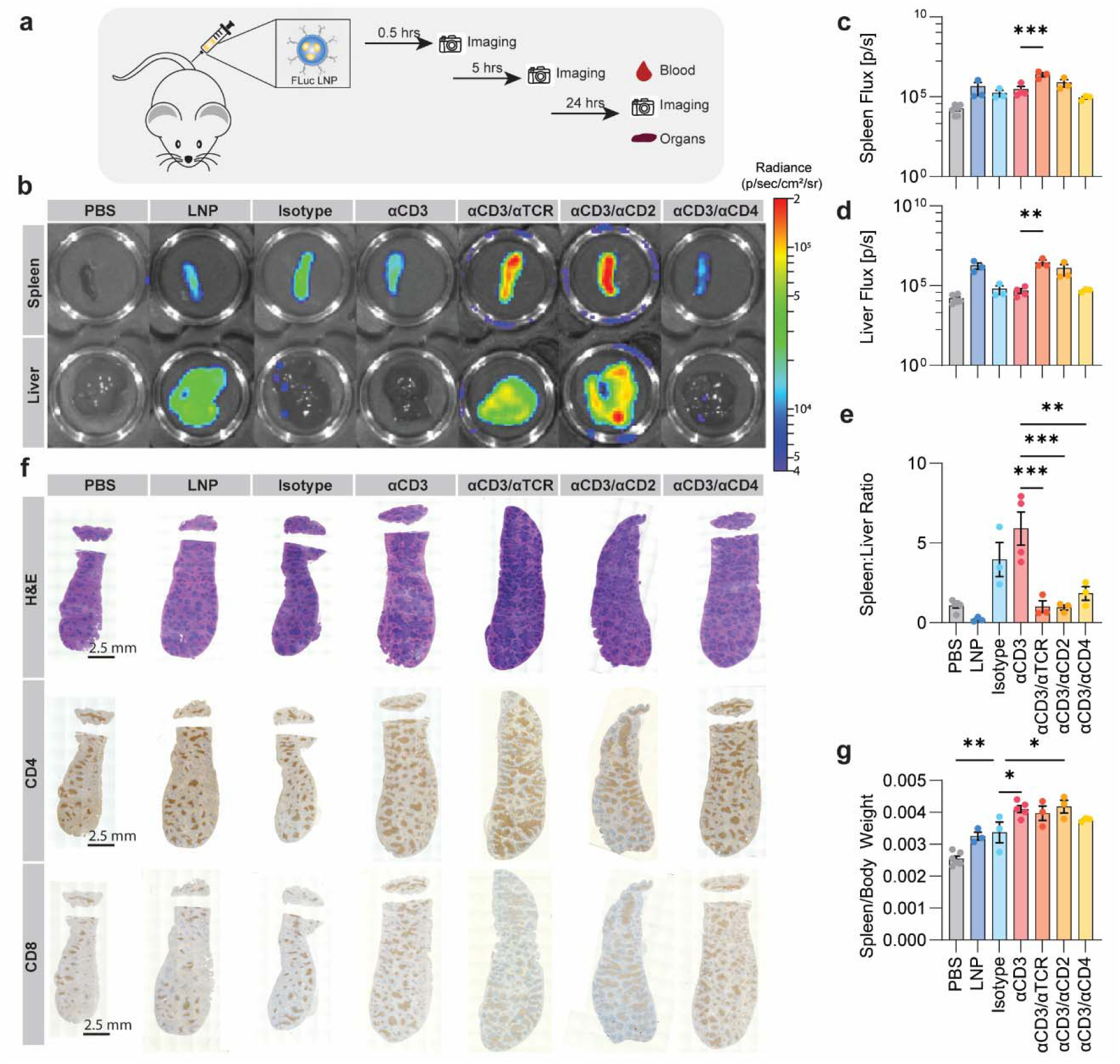
tLNPs facilitate transfection of the spleen. (a) Schematic of workflow for studying biodistribution of tLNPs. (b) Representative IVIS images of organs harvested from mice 24 h after intravenous injection of LNPs containing Fluc mRNA at a dose of 0.6 mg/kg body weight. Measurements of luminescence for the spleen (c) and liver (d) as regions of interest on the IVIS (n ≥ 3 biological replicates, error bars = standard error of the mean, outlier removed using ROUT Q = 1%). (e) Ratio of the luminescence for the spleen and liver of each mouse. Statistical analysis included ordinary one-way ANOVA with Dunnett**’**s multiple comparisons test, **p<0.01 and ***p<0.001 as compared to αCD3-LNP. (f) Histological sections of the spleen harvested 24 h after intravenous injection stained with H&E, αCD4, and αCD8. Scale bars are 2.5 mm. (g) Spleen weight normalized to mouse body weight 24 h after treatment. Statistical analysis included ordinary one-way ANOVA with Dunnett**’**s multiple comparisons test *p<0.05 and **p<0.01 as compared to isotype-LNP.

We first evaluated the organ specific transfection of tLNPs, focusing on the liver and spleen, but also evaluated other organs (*SI Appendix*, Fig. S9). The images of luminescence from the spleen and liver revealed differences in the biodistribution of the tLNPs as a function of their surface targeting (Fig. 4*B*). After selecting a consistent region of interest and normalizing to the background, the flux was calculated revealing that αCD3/αTCR-LNPs had the highest transfection in the liver and in the spleen (Fig. 4*C*). tLNPs shifted transfection towards the spleen from the liver relative to untargeted LNPs. The highest spleen to liver luminescence ratio occurred upon administration of the αCD3-LNPs. This ratio declined for LNPs targeted with αCD3/TCR and αCD3/αCD2-LNP because although there was higher luminescence observed in the spleen, there was also much greater luminescence in the liver (Fig. 4*D*). αCD3/αCD4-LNP had the lowest luminescence observed in the spleen of all tLNP groups possibly due to targeting of a subset of T cells rather than the entire T cell population. In contrast, untargeted LNPs predominantly transfected the liver, leading to the lowest spleen: liver luminescence ratio of all formulations (Fig. 4*E*). This aligns with previous work which has shown that Dlin-MC3-DMA LNP**’**s tend to accumulate in the liver (26, 30, 31).

At 24 h the physical characteristics of the spleen were analyzed for spatial distribution of T cells and physical enlargement because the white pulp of the spleen is a major site for T cell activation and proliferation (32, 33). Histological sections of the spleen were stained using hematoxylin and eosin (H&E), and CD4 and CD8 markers (Fig. 4*F*). H&E revealed expanded of areas of white pulp (dark purple) for αCD3-LNPs and bispecific LNPs as compared to the control groups (Fig. 4*F*). The CD4^+^ and CD8^+^ cells appeared to remain concentrated in the periarterial lymphatic sheath (PALS) for all groups however there was a remarkable expansion in these populations in αCD3-LNP and bispecific groups in comparison to the control groups indicating that these cells were encountering T cell specific antibodies resulting in increased T cell production within the spleen (Fig. 4*F*). T cell proliferation generally follows T cell activation indicating an immune response triggered by tLNPs. This proliferation was not seen in Iso-LNP treatment indicating that the proliferation is due to the T cell specific ligands and not due to the LNP, mRNA, DNA-tethers, or overall antibody structure. In addition to this visual increase in splenic T cells, the overall spleen mass was also found to increase with the addition of T cell specific antibodies as compared to the unmodified LNP group indicating enlargement attributed to αCD3 and bispecific surface modification (Fig. 4*F*). Overall, this shows that antibody surface modification can impact LNP tropism, however other factors such as LNP lipid composition should also be considered.

### Select bispecific LNP formulations enhance T cell transfection and diminish CD3 depletion *in vivo*

Next, we evaluated the cell-specific transfection of tLNPs *in vivo*. Here, we analyzed T cells (CD3^+^, CD4^+^, and CD8^+^), B cells (CD19^+^), and monocytes (CD11b^+^) present in the blood, spleen, and lymph nodes 6 h after intravenous injection (Fig. 5*A*). LNP formulations were assembled encapsulating mRNA encoding mCherry and the % of mCherry^+^ cells in the aforementioned regions were analyzed using flow cytometry (*SI Appendix*, Fig. S11). In the blood, we observed all bispecific LNP formulations improved total T cell transfection (CD3^+^) relative to αCD3 LNP and all LNP controls. The αCD3/αTCR-LNP had the highest % of mCherry^+^ cells in combined CD4^+^ and CD8^+^ T cells in the blood with ∼10% transfection efficiency relative to the monospecific αCD3 LNP (3% transfection) and LNP controls (< than 2 % transfection). The improved transfection of T cells with bispecific LNPs may be due to the capacity of these LNPs to engage with the T cell receptor complex at two different epitopes causing an increase in receptor mediated endocytosis (Fig. 5*B*). In blood resident monocytes, we observed bispecific LNPs also resulted in increased transfection relative to the monospecific αCD3 LNP and control LNPs. We hypothesize this enhanced transfection of monocytes may be due to the presence of additional Fc regions on the bispecific LNP surface resulting in increased engagement with scavenger receptors on monocytes (34). To mitigate this uptake, it is advisable to use antibody fragments (i.e. nanobodies or single-chain variable fragments) rather than full length IgGs in downstream tLNP formulations. In the spleen, DNA-modified LNPs led to higher transfection efficiencies than the unmodified LNP, suggesting that the modification chemistry has some capacity to increase splenic T cell transfection. (Fig. 5*D*). Transfection of spleen resident monocytes remained low for all groups (Fig. 5*E*). Overall, the enhanced transfection of T cells at levels as high as 10% are consistent with levels needed for *in vivo* gene delivery approaches like CAR T therapies and support the capacity of DNA-anchoring to assemble tLNPs with therapeutic potential (8).

**Figure 5.**
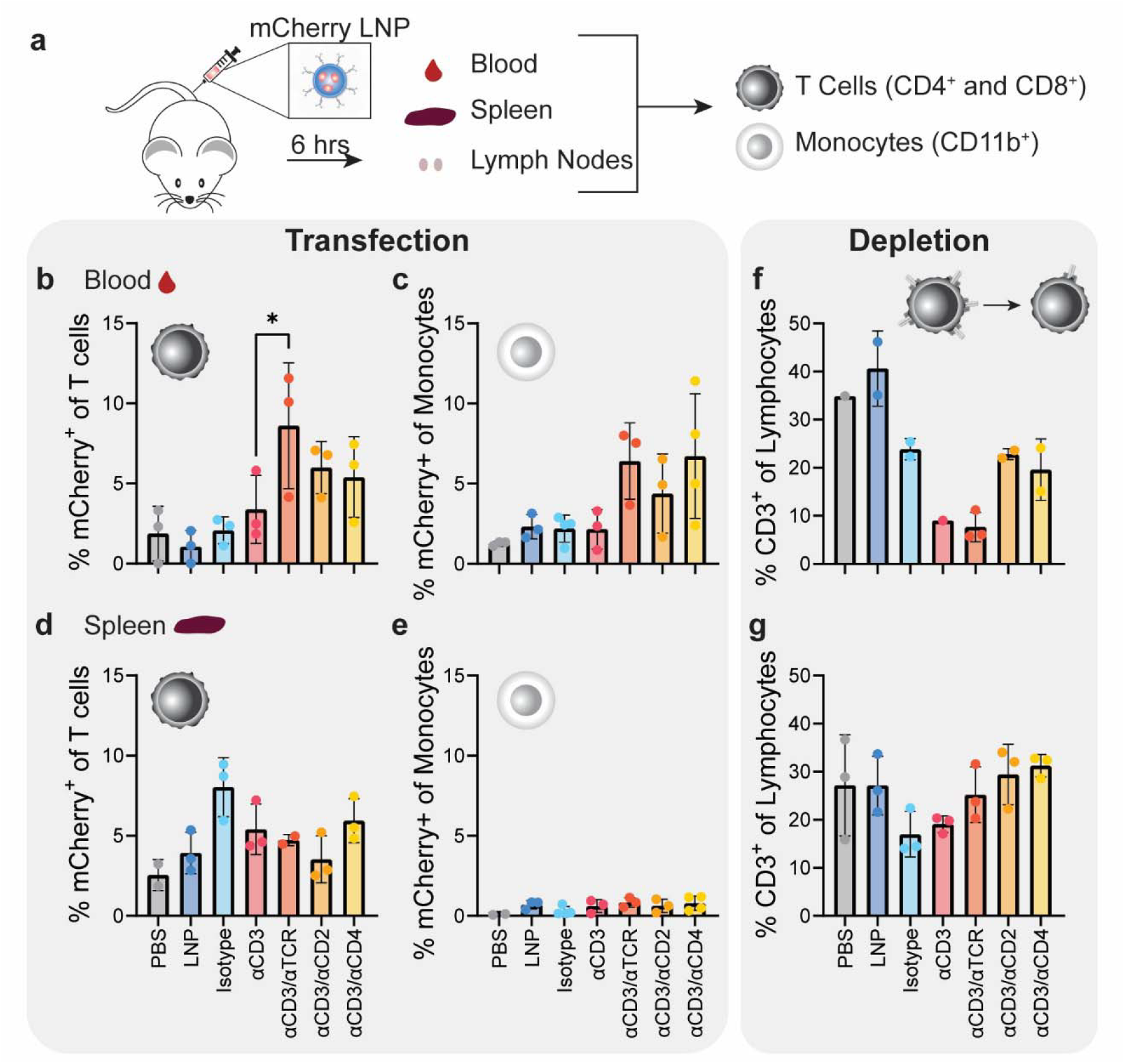
Select LNP formulations enhance T cell transfection and diminish depletion. (a) Schematic of the workflow for *in vivo* cell transfection study. Flow cytometry data showing the percent of T cells and monocytes expressing mCherry in the blood (b-c) and spleen (d-e) and the percent of CD3^+^ cells in the blood (f) and spleen (g). Statistical analysis included ordinary one-way ANOVA multiple comparisons test, *p<0.05 and **p<0.01. (n = 3 biological replicates, error bars = standard error of the mean, samples with low cell counts were omitted from data sets).

In the blood, as we previously observed in PBMCs *in vitro*, there was a depletion of CD3^+^ cells for all tLNPs groups (Fig. 5*F*). Depletion of the CD3 surface marker correlates with the internalization of the T cell receptor complex, the presentation of which is key for proper T cell activation and mounting a functional immune response. The αCD3-tLNPs and αCD3/αTCR-tLNPs resulted in the greatest depletion of CD3^+^ cells in the blood. αCD3/αCD2 and αCD3/αCD4-tLNPs, however, resulted in less depletion of the T cell receptor complex while still increasing transfection of T cells in the blood as compared to the αCD3-tLNPs. A similar trend of CD3 depletion with αCD3-tLNPs was also seen in the spleen (Fig. 5*G*). These findings indicate that the choice of tLNP antibodies not only influences TCR expression levels post-transfection but may also impact downstream TCR signaling and functional activation.

## Discussion

Using DNA-tethered conjugation of full-length commercial antibodies to Dlin-MC3-DMA LNPs we demonstrate that tLNPs increase the efficiency of *in vivo* T cell transfection through specific antibody–receptor interactions. Specifically, we found that bispecific LNPs increase T cell transfection over single antibody formulations in both *ex vivo* studies with PBMCs and *in vivo* in the blood, showing that co-targeting of receptors can be used to improve cell-selective mRNA delivery.

This work highlights the significance of targeting the T cell receptor (TCR) complex as a strategic approach to improving the efficiency of T cell transfection. The TCR complex is generally stably expressed and rapidly recycled in resting T cells (35). This receptor internalization allows for therapeutic nanoparticles, like tLNPs, to enter T cells via binding the TCR complex receptors and overcome the T cells natural resistance to particle uptake. Bispecific LNPs, which included TCR or CD3 targeting antibodies, were more potent transfection agents than LNPs which did not target the T cell receptor. The bispecific αCD3/αTCR-tLNP achieved the highest transfection rates in circulating T cells (∼10%) (Fig. 5*A*) and highest splenic transfection (Fig. 4*C*). This suggests that targeting two epitopes on the same receptor complex creates a synergistic effect, which can also influence where nanoparticles distribute in the body. This approach may come with drawbacks, however, as indicated by the increased CD3^+^ depletion observed when T cells are engaged only through the TCR complex and without co-stimulation. In such cases, T cells with low CD3^+^ expression can become functionally unresponsive although some reports have also shown the ability of T cells to recover CD3^+^ expression after complete clearance of αCD3 from systemic circulation (9, 36). Although outside the scope of the current project, further work should be done to understand how CD3 depletion may affect subsequent T cell functional activation. To circumvent CD3^+^ depletion while maintaining transfection and cell-specific delivery that results from targeting the TCR complex, bispecific LNPs including a costimulatory ligand may offer a more effective therapeutic strategy for *in vivo* gene delivery.

In our study, bispecific tLNPs, that included co-stimulatory molecules such as αCD3/αCD2-LNPs and αCD3/αCD4-LNPs, exhibited increased transfection in the blood compared to αCD3-LNPs. They also increased transfection in a specific T cell subset, in this case CD4^+^ helper T cells. Notably CD3 expression increased in these bispecific groups compared to αCD3 alone, mitigating excessive receptor depletion. In previous work, bispecific tLNPs designed to simultaneously engage CD3 and a co-stimulatory receptor—examples include CD3/CD28 (13) and CD3/CD7 (16)—exhibited enhanced activity relative to single-targeted counterparts, a trend mirrored in our results. The choice of costimulatory molecules appears to influence both cell-specific mRNA delivery as well as cell activation capacity.

Transfection of circulating T cells is important for *in vivo* cellular engineering and immune modulation and our transfection of ∼10% of circulating T cells falls within the desired range needed to achieve a therapeutic effect for *in vivo* CAR T generation and immunomodulation. More importantly, the DNA-tethering approach streamlines conjugation without the limitations of traditional chemical methods, enabling rapid, high-throughput *in vivo* screening of new bispecific or multivalent formulations. We anticipate the ease of assembling multivalent tLNPs and nanoparticles with this approach will allow for rapid selection of nanoparticle targeting ligands that are disease and target cell specific. Furthermore, the modularity of this approach allows facile exchange of targeting molecules, stoichiometry of multiple antibodies, and precise control of tether length. This physicochemical control of tLNPs supports the exploration of diverse receptor combinations to optimize specificity, efficacy, and biodistribution for future antibody–LNP therapeutics.

## Materials and Methods

### Materials

The following chemicals were used as received: Acetic acid, glacial, 99+% (AA33252AK, Thermo Scientific Chemicals), chloroform, stabilized with amylene for HPLC (C297-4, Fisher Chemical), ethanol, absolute (200 proof), molecular biology grade (BP28184, Fisher BioReagents), methanol (A452SK-4, Fisher Chemical), and EDTA (AM9260G, Invitrogen). 1,2-distearoyl-sn-glycero-3-phosphoethanolamine-N-[azido(polyethylene glycol)-2000] (ammonium salt) (DSPE-PEG(2000) Azide, 880228), cholesterol (ovine, 700000), 1,2-distearoyl-sn-glycero-3-phosphoethanolamine-N-[methoxy(polyethylene glycol)-2000] (ammonium salt) (DSPE-PEG2000, 880120), 1.2-distearoyl-sn-glycero-3-phosphocholine (DSPC, 850365), 1,2-distearoyl-sn-glycero-3-phosphoethanolamine-N-[amino(polyethylene glycol)-2000]-N-(Cyanine 7) (DSPE PEG(2000)-N-Cy7, 810892), and 1,2-dioleoyl-sn-glycero-3-phosphoethanolamine-N-(lissamine rhodamine B sulfonyl) (ammonium salt) (18:1 Liss Rhod PE, 810150) were purchased from Avanti Polar Lipids. D-Lin-MC3-DMA (MC3, BP-25497) was purchased from BroadPharm. SYBR gold nucleic acid gel stain (S11494), Quant-iT OliGreen ssDNA Reagent and Kit (O11492), and Quant-it RiboGreen RNA Assay Kit (R11490) were purchased from Invitrogen. 2-mercaptoethanol (1610710), QC colloidal Coomassie (1610803), precision plus protein dual color standards (1610374), and 4-20% polyacrylamide gradient gel (4561096) were purchased from Bio-Rad. Dulbecco**’**s phosphate-buffered saline, no calcium, no magnesium (dPBS, 14190), RPMI 1640 Medium (11875093), penicillin streptomycin (pennstrep, 15-140-122) and fetal bovine serum, heat inactivated (A5669801) were purchased from Gibco. 1x red blood cell lysis buffer (00-4333-57) was purchased from eBioscience. Sepharose 4B (4B200) and Sepharose CL-2B (CL2B300) were purchased from Sigma-Aldrich. OYo-Link custom oligo reagent (5**’** TTA ATC ACC CTC GCG CAC TAC 3**’**, AT1002) for modification for general antibody modification and for mouse IgG1 modification was purchased from AlphaThera. 5**’** -/5ATTO633N/TTA ATC ACC CTC GCG CAC TAC 3**’** and 5**’** /5DBCON/CTA GTG CGC GAG GGT GA 3**’** were purchased from IDT. For surface modification of LNPs for *in vitro* studies, Alexa Fluor 647 anti-human CD28 antibody (302953), Ultra-Leaf purified anti-human CD3 antibody (300438), purified anti-human TCR α/β antibody (306702), purified anti-human CD2 antibody (300202), Ultra-LEAF purified anti-human CD4 antibody (300569), purified anti-human CD5 antibody (300602), purified anti-human CD8 antibody (344702), Ultra-LEAF purified anti-human CD28 antibody (302934), and purified moused IgG1, κ isotype control antibody (400102) were purchased from Biolegend. Purified mouse anti-human CD6 (555356) and purified mouse anti-human CD7 (555359) were purchased from BD Biosciences. For surface modification of LNPs for *in vivo* studies, CD3e monoclonal antibody (16-0031-82), TCR beta monoclonal antibody (16-5961-82), CD2 recombinant rabbit monoclonal antibody (18056481), CD4 monoclonal antibody (16-0041-82), and rat IgG2b κ isotype control (16-4888-81) were purchased from Invitrogen. ONE-Glo + Tox Luciferase Reporter and Cell Viability Assay (E7120) and VivoGlo Luciferin (P1043) were purchased from Promega. mCherry mRNA (L-7203) and luciferase mRNA (L-7602) were purchased from TriLink. Saline sodium chloride 0.9% (04888010) was purchased from Hospira. For flow cytometry studies on human cells, Zombie UV Fixable Viability Kit (423107), Human TruStain Fc receptor blocking solution (422302), APC-Cyanine7 anti-human CD3 antibody (344818), CD4 Pac Blue 317429 Pacific Blue anti-human CD4 antibody (317429), CD8 FITC anti-human CD8a antibody (300906), CD14 APC anti-human CD14 antibody (325607), and CD19 PE/Cyanine7 anti-human CD19 antibody (302216) were purchased from Biolegend. For flow cytometry studies on mouse cells, TruStain FcX PLUS (anti-mouse CD16/32, 156603), Spark Red 718 anti-mouse CD3 antibody (100282), Alexa Fluor 647 anti-mouse CD4 antibody (100426), Alexa Fluor 488 anti-mouse CD8a antibody (100726), Brilliant Violet 421 anti-mouse CD19 antibody (115537), and PE anti-mouse/human CD11b antibody (101207) were purchased from Biolegend.

### Methods

#### Antibody Modification and Characterization

Full length IgG antibodies were conjugated to single stranded DNA using light activated site-specific conjugation (27). oYo-Link mIgG1 was used for all IgG1 antibodies with the mouse host species. All other antibodies were modified with oYo-Link.100ug of oYo-Link custom oligo reagent (5**’** TTA ATC ACC CTC GCG CAC TAC 3**’**) was rehydrated in 100 uL of ultrapure water. 1 uL of oYo-Link reagent was mixed with 1 ug of antibody and then placed under ∼365 nm UV light (AT8001-D, AlphaThera) for 2 hours.

A reducing SDS-PAGE gel was used to analyze antibody modification. Prepare samples at 0.5 uM antibody in with 10 % volume of 2-mercaptoethanol. The samples were heated for 10 min at 90°C. 10 uL of each sample was loaded into a 4-20% polyacrylamide gradient gel. 3 uL of the protein ladder was loaded into well 1. The gels ran for ∼1 hour at 120 volts. The gel was incubated in 1x SYBR Gold for 10 minutes before imaging using AzureRed fluorescent imaging (Azure 200, Azure Biosystems). The gel was then fixed in a solution of 50% v/v methanol and 1% v/v acetic acid for 15 minutes before incubating in colloidal Coomassie for 2 hours. Gels were imaged using Coomassie blue settings (Azure Biosystems).

#### DNA-Lipid Synthesis

As previously described by Banga et al. (25), DNA conjugated lipids were synthesized using copper-free click chemistry. Briefly, the DBCO-terminated oligonucleotide (5**’** /5DBCON/CTA GTG CGC GAG GGT GA 3**’**) was dissolved in 100 uL ultrapure distilled water at a concentration of 100 uM. In a separate glass vial, 100 nmol (28 uL at 10 mg/mL) of DSPE-PEG(2000) Azide in chloroform was dried under inert nitrogen gas and then dissolved in 100 uL ethanol. The aqueous oligonucleotide solution was then added to the lipid solution and the mixture was left to shake on the benchtop at room temperature. The next day the contents of the vial were dried under inert nitrogen gas. The dried material was resuspended in 300 uL ultrapure distilled water. Excess lipid was purified using chloroform extraction. Briefly, 300 uL of chloroform was added to the solution and mixed through gentle inversion before spinning for 5 min at 16,000 g and removing the top aqueous layer. This chloroform wash was repeated 3 times. Concentration of DNA-lipid was determined using Quant-iT OliGreen ssDNA Reagent and Kit according to the manufacturer**’**s instructions. The molecular weight of the DNA-lipid was confirmed using mass spectrometry at the IMSERC Mass Spectrometry Facility.

#### LNP Formulation and Characterization

LNPs were prepared at the molar ratio of 50:13:35:2 of MC3:DSPC:cholesterol:DSPE-PEG2000. For dyed LNPs, the 2 mol% DSPE-PEG2000 was replaced with 1.7 mol% DSPE-PEG2000 and 0.3 mol% DSPE PEG(2000)-N-Cy7 or 18:1 Liss Rhod PE. Lipids were dissolved in ethanol to form the ethanol phase. 30 uL mRNA at a concentration of 1 mg/mL or 30 uL PBS for empty LNPs, 22.5 uL citrate buffer at a concentration of 100 mM, and 172.5 uL ultrapure water were combined in a glass vial to form the aqueous phase. The aqueous phase was mixed using a stir bar and stir plate (Fisher Scientific) at 700 rpm while the ethanol phase was added dropwise to the aqueous phase. LNPs were dialyzed overnight at 4°C

Encapsulation efficiency was measured for LNPs using Quant-it RiboGreen RNA Assay Kit according to the manufacturer**’**s instructions. LNP size and polydispersity were determined using dynamic light scattering (Malvern Zetasizer Ultra). 30 uM LNP were read for each experiment, and each replicate is the average of 3 reads of a single sample.

#### Ab-LNP Conjugation and Characterization

LNPs were modified through incubation of oYo-Link modified antibody at a ratio of 0.02 uM/1mM LNP lipid, DNA-lipid at a ratio of 1.8 uM/1mM LNP lipid, and LNPs at 37°C for 1 hour. For bispecific LNPs, modified antibodies were added at a 1:1 ratio with the ratio of 1.8 uM/1mM LNP lipid remaining the same. For transfection of human cells *in vitro* (Jukat cells and PBMCs), LNPs were modified with anti-human CD3, anti-human TCR, anti-human CD2, anti-human CD4, anti-human CD5, anti-human CD6, anti-human CD7, anti-human CD8, anti-human CD28, and IgG1 κ isotype control. For transfection of mouse cells *in vivo*, CD3e monoclonal antibody, TCR beta monoclonal antibody, CD2 recombinant rabbit monoclonal antibody, CD4 monoclonal antibody, and rat IgG2b κ isotype control were used.

Size exclusion chromatography was used to determine DNA-lipid insertion into and antibody conjugation to LNPs. Following LNP modification with DNA-lipid, a complementary fluorescent ssDNA (5**’** -/5ATTO633N/TTA ATC ACC CTC GCG CAC TAC 3**’**) was incubated in excess with DNA-LNPs at 37°C for 1 hour. DNA-LNP samples were loaded into a column packed with Sepharose 4B and fractions were collected in a 96 well plate using a Gilson FC204 Fraction Collector. Plates were then read on a Molecular Devices Spectra Max iD3 plate reader for ATTO 633 (ex 620 nm/em 660 nm) and Liss Rhod (ex 550 nm/em 590 nm). For Ab-LNPs, an Alexa Fluor 647 anti-human CD28 antibody was conjugated to the surface. Ab-LNPs were loaded into a column packed with Sepharose CL-2B. Fractions were collected as previously described. Plates were read for CD28 antibody (ex 630/em 670) and Cy7 (ex 745 nm/em 785 nm). To determine the concentration of DNA or antibody conjugated to the LNP, the measured fluorescence in LNP (void volume) fractions was compared to a standard curve for each fluorophore. LNP concentrations were determined using nanoparticle tracking analysis. Samples with a lipid concentration of 0.15 mM were analyzed using NanoSight300 (Malvern Instruments).

#### Cell Culture

Jurkat cells were purchased from ATCC (TIB-152). PBMC**’**s were generously gifted from the Choi Lab (Northwestern Medicine) or purchased from ATCC (PCS-800-011). Cells were cultured in RPMI 1640 Medium supplemented with 10% FBS and 1% pennstrep.

#### In Vitro Jurkat Cell Luciferase and Toxicity Assay

Jukat cells were plated in triplicate at 100,000 cells per 50 uL supplemented RPMI. 50 uL LNPs suspended in PBS (containing 750 ng luciferase mRNA) were added to each well. After 24 hours incubation at 37°C, ONE-Glo + Tox Luciferase Reporter and Cell Viability Assay was used to determine cell luminescence and LNP toxicity according to the manufacturer**’**s instructions. Briefly, 20 uL 5X CellTiter-Fluor Reagent to each well and mix using orbital shaking for ∼30 seconds. After 30-minute incubation at 37°C, fluorescence was measured on a Molecular Devices Spectra Max iD3 plate reader (ex 465 nm/em 505 nm). Next, 100 uL ONE-Glo Reagent was added to each well and was incubated for 3 minutes at room temperature before luminescence was measured on a Molecular Devices Spectra Max iD3 plate reader.

#### In vitro PBMC Transfection using mCherry mRNA

PBMC**’**s were thawed and plated in a round bottom 96 well plate at 300,000 cells per 100 uL supplemented RPMI. 100 uL LNPs suspended in PBS containing 2000 ng mCherry mRNA were added to each well. After 24 hours incubation at 37°C, cells were suspended in dPBS and stained with Zombie UV Fixable Viability Kit according to the manufacturer**’**s instructions. Cells were then suspended in 100 uL flow buffer consisting of dPBS supplemented with 1% FBS and 2 mM EDTA and blocked using Human TruStain Fc receptor blocking solution. Cells were stained with APC-Cyanine7 anti-human CD3 antibody, Pacific Blue anti-human CD4 antibody, FITC anti-human CD8a antibody, APC anti-human CD14 antibody, and PE/Cyanine7 anti-human CD19 antibody for 30 minutes at 4°C. Stained cells were analyzed on LSR Fortessa 2 Analyzer (BD Biosciences) and data sets were analyzed on FlowJo 10.10.0 (BD Biosciences).

#### Animals

All animal studies were conducted according to the guidelines for the Care and Use of Laboratory Animals from the National Institutes of Health (37) and all animal work was performed under protocols approved by the Institutional Animal Care and Use Committee (IACUC). Male BALB/c mice ∼10-12 weeks in age were purchased from Jackson Labs used in all animal studies. Mice were placed on a low-fluorescent diet several days prior to all experiments.

#### In Vivo Biodistribution using Luciferase mRNA

LNPs containing a Cy7 dye and luciferase mRNA were given via tail vein injection at a dose of ∼0.6 mg Fluc mRNA/kg body weight. *In vivo* images were taken using IVIS Spectrum (PerkinElmer, Waltham, MA) for the fluorescence intensity of Cy7 (ex 745 nm/em 780 nm) and bioluminescence of Fluc at 0.5, 5, and 24 hours. D-luciferin in saline was given at 75 mg/kg body weight via intraperitoneal injection 10 minutes prior to imaging. Blood was collected into EDTA-treated tubes (365974, Fisher Scientific) using retroorbital bleeding at 0.5 and 5 hours. 24 hours following tail vein injection, mice were euthanized, and blood was collected using cardiac puncture. The left and right inguinal lymph nodes, spleen, liver, heart, kidneys, lungs, skin, and leg were imaged *ex vivo* for Cy7 fluorescence and bioluminescence of luciferase using the IVIS Spectrum (PerkinElmer) and the mass of each organ was recorded.

#### In Vivo Delivery using mCherry mRNA

LNPs containing a Cy7 dye and mCherry mRNA were given via tail vein injection at a dose of ∼0.6 mg Fluc mRNA/kg body weight.6 hours after LNP injection, blood was collected using heart puncture into EDTA-treated tubes. Red blood cells were lysed using 1x red blood cell lysis buffer. The spleen and left and right inguinal lymph nodes were collected and pressed through a 40-um cell strainer (08-771-1, FisherScientific) to create a single cell suspension. Cell suspensions were stained using Zombie UV Fixable Viability Kit according to the manufacturer**’**s instructions. Cells were then suspended in 100 uL flow buffer and blocked using TruStain FcX PLUS. Cells were stained with Spark Red 718 anti-mouse CD3 antibody, Alexa Fluor 647 anti-mouse CD4 antibody, Alexa Fluor 488 anti-mouse CD8a antibody, Brilliant Violet 421 anti-mouse CD19 antibody, and PE anti-mouse/human CD11b antibody.

#### Statistics

Statistical analyses were performed using Prism 10.4.0 software (Graph-Pad Software Inc.) Statistical tests are reported within each figure caption. P-values of <0.05 were considered significant and were adjusted for multiple comparisons. Data are expressed as mean ± SEM, unless otherwise specified.

## Acknowledgments

This work was primarily supported by the Northwestern McCormick Research Catalyst Award (N.P.K.). This work was partially supported by NSF (DMR-2145050) to N.P.K. M.D.K was supported by National Institutes of Health Training Grant (T32-EB031527) from the Northwestern University**’**s Regenerative Engineering Training Program. T.Q.V. and L.C. were supported by the National Institutes of Health Training Grant (T32GM008449) through Northwestern University**’**s Biotechnology Training Program. C.S. was supported by an NSF Graduate Research Fellowship. This work made use of the Keck-II and BioCryo facilities of Northwestern University**’**s NUANCE Center, which has received support from the SHyNE Resource (NSF ECCS-2025633), and Northwestern**’**s MRSEC program (DMR-2308691). This work was also supported in part by the Northwestern University Flow Cytometry Core Facility supported by the Cancer Center Support Grant (NCI 5P30CA060553). The authors thank Kim Cardenas for her assistance in planning flow cytometry studies. Primary T cells were generously gifted by the laboratory of Prof. Jaehyuk Choi. We thank the Ameer Laboratory for use of their transmitted light microscope (Carl Zeiss AG, Germany) for imaging of histological sections. Mass spectrometry was performed at the Integrated Molecular Structure Education and Research Center at Northwestern. Histological samples were processed by the Mouse Histology and Phenotyping Laboratory at the Robert H. Lurie Comprehensive Cancer Center (NCI P30-CA060553). Animal studies were performed by the Northwestern University Developmental Therapeutics Core and IVIS imaging work was performed at the Northwestern University Center for Advanced Molecular Imaging (Evanston), both supported by NCI CCSG P30 CA060553 awarded to the Robert H Lurie Comprehensive Cancer Center. The authors thank Nayereh Ghoreishi–Haack for her assistance planning the animal studies, and Elizabeth Dempsey for her expertise in both planning and executing the animal studies.

