## Supplemental Information for "DNA-Directed Assembly of Multivalent Lipid Nanoparticles for Targeted T Cell Gene Delivery"

*Neha P. Kamat

**Figures**

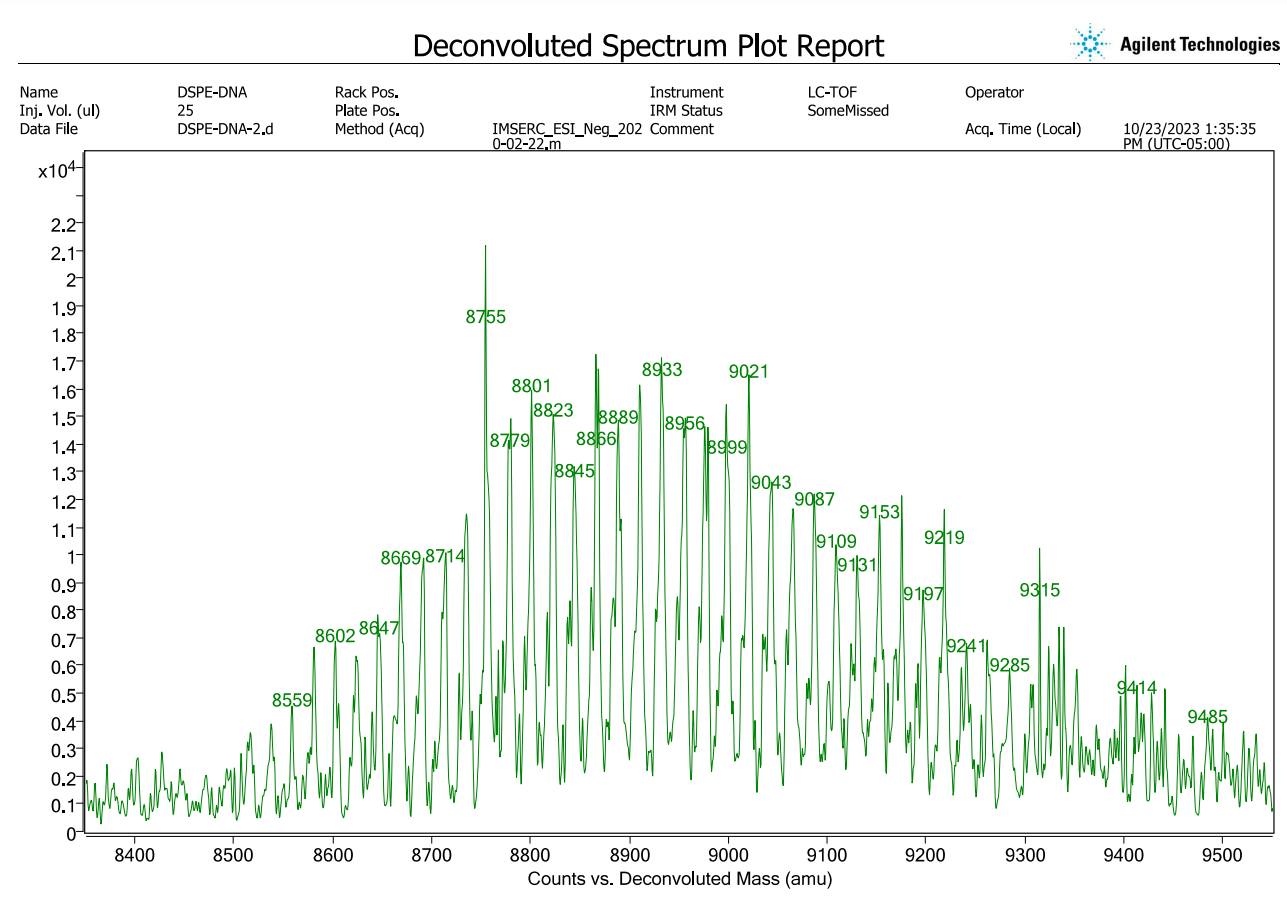

Fig. S1. Mass spectrometry of DNA-conjugated lipid. Mass spectrometry showing a mass of approximately 8,700 amu equivalent to the addition of the DNA molecular weight (5,827 amu) and the lipid molecular weight (2,816.52 amu).

**Fig. S2. Validation of antibody modification.** A protein gel (left) and DNA stain (right) were used to confirm antibody modification with DNA (AlphaThera).

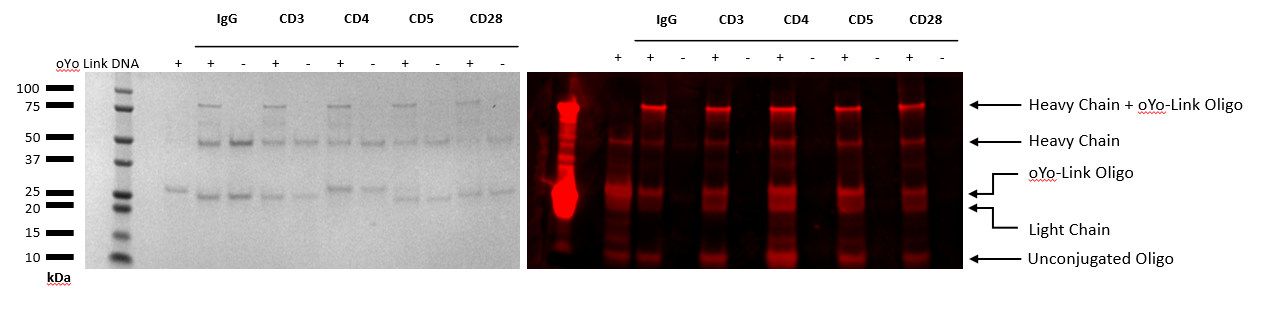

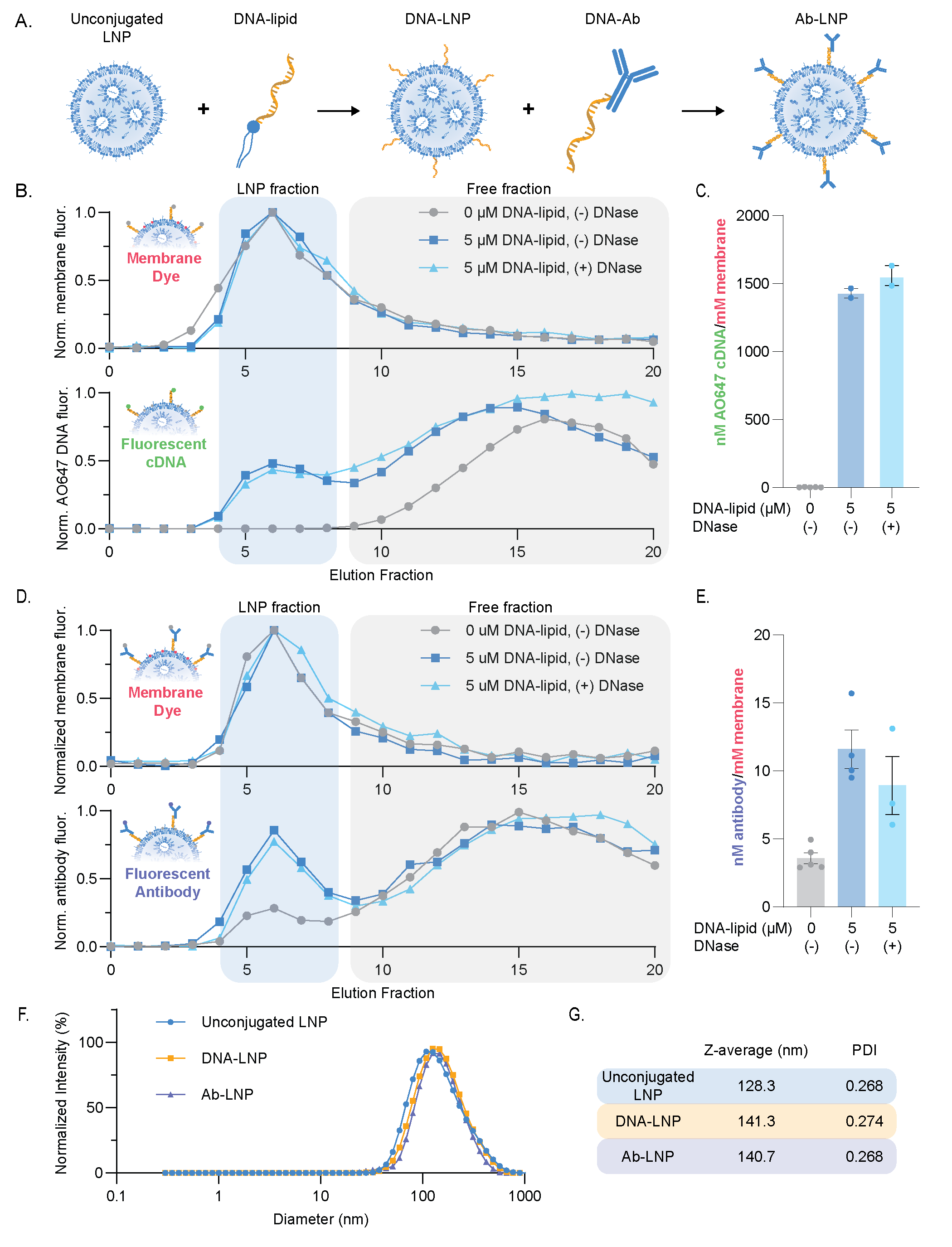

**Fig. S3. Validation of particle modification with DNA/antibody using size exclusion chromatography.** Particles containing a fluorescent rhodamine dye (a) were conjugated to a fluorescent DNA probe (upper panel) or Alexa 647 CD28 antibody (lower panel) and run on size exclusion chromatography. The coelution of the DNA or antibody with the LNPs indicated successful conjugation.

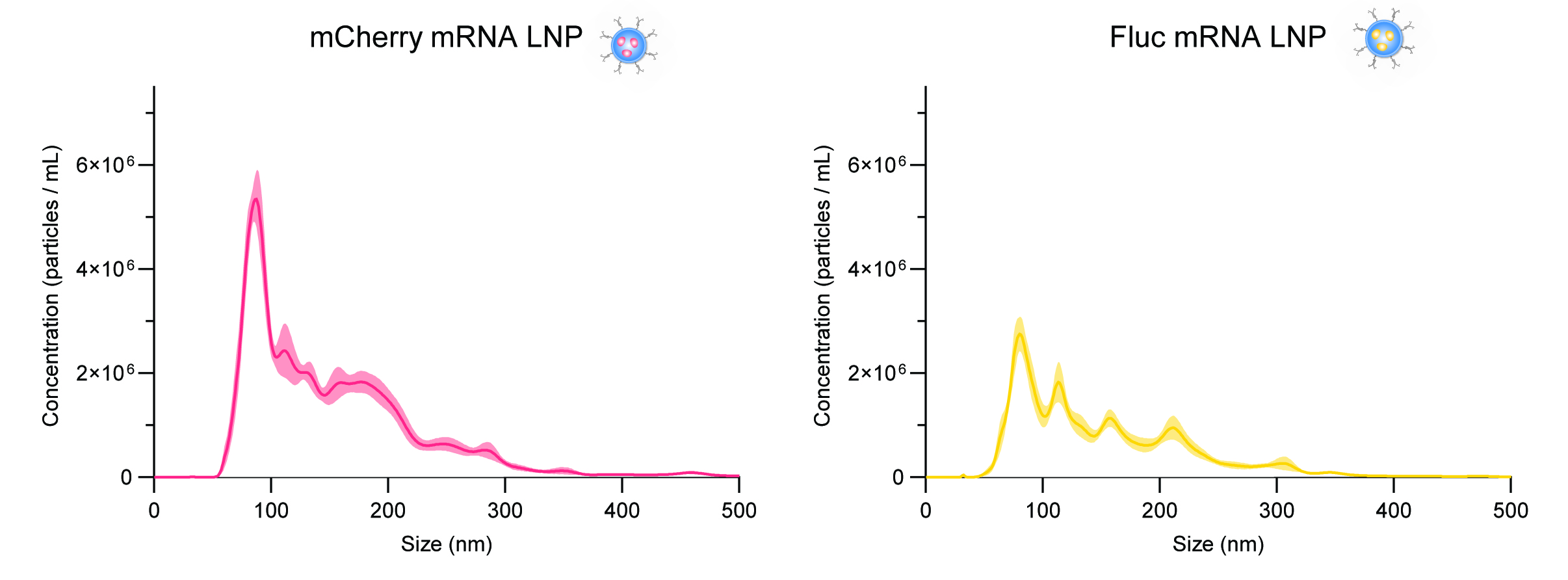

**Fig. S4. Nanoparticle tracking analysis showing distribution of size of LNPs.** The size and concentration of mCherry and Fluc LNPs was determined using particle tracking analysis.

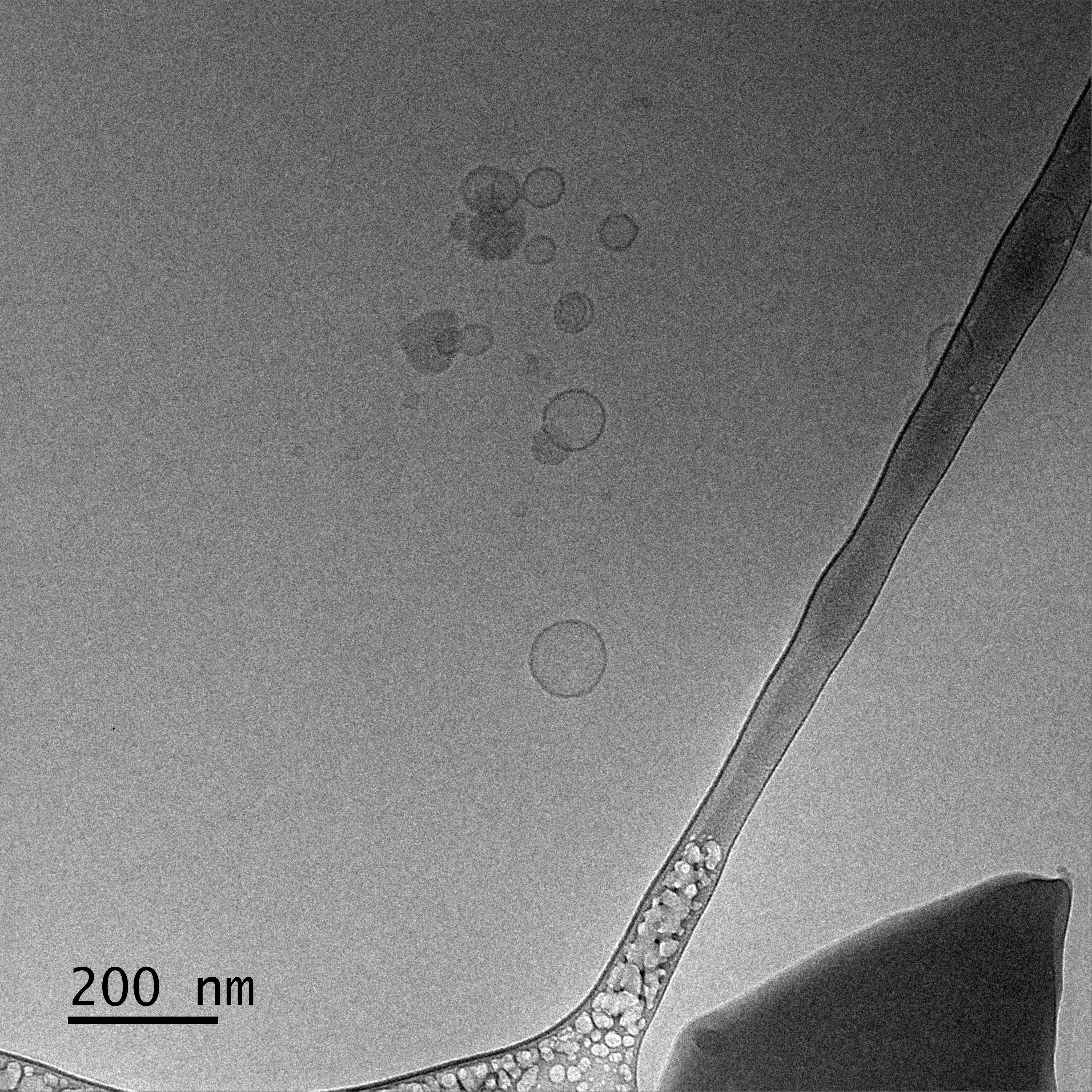

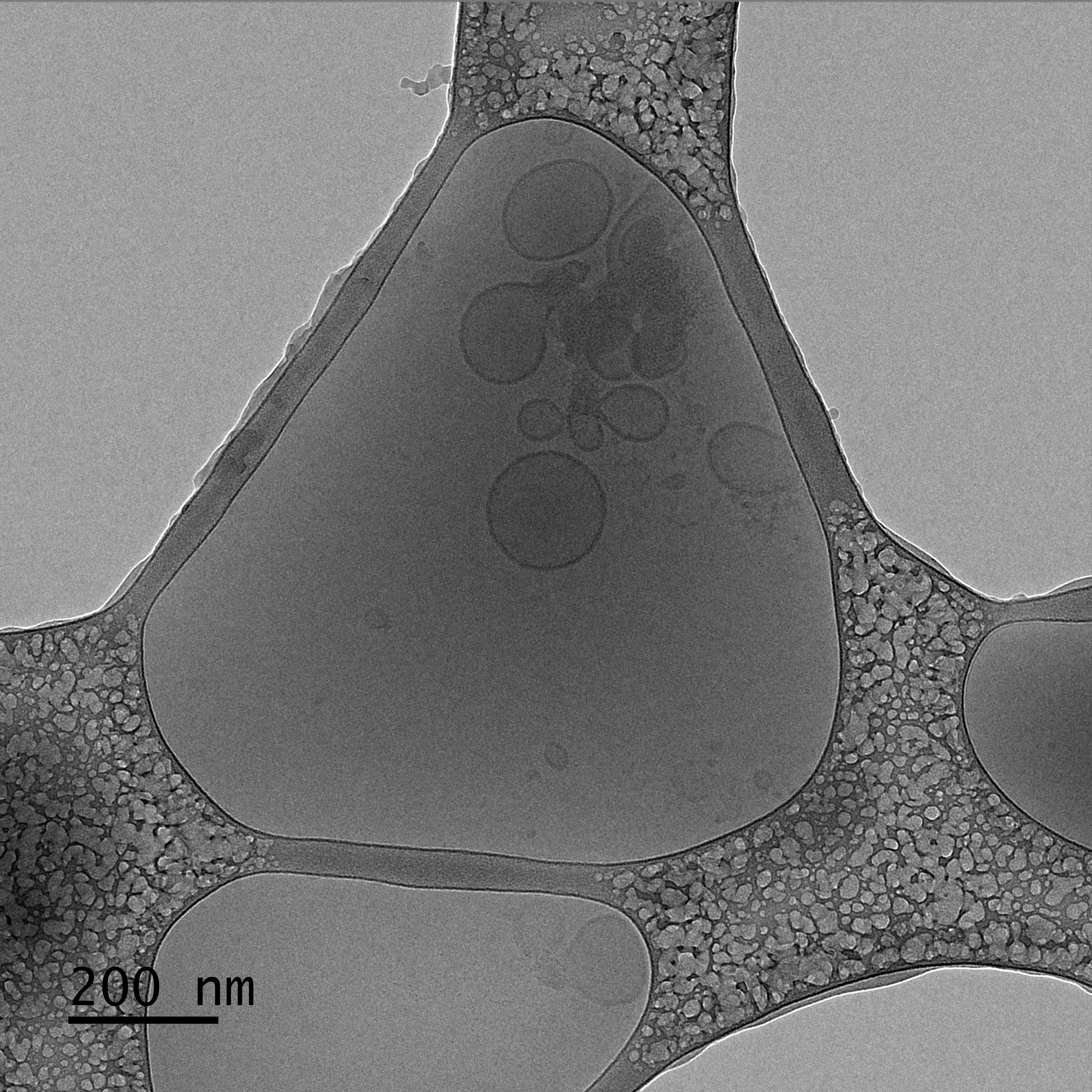

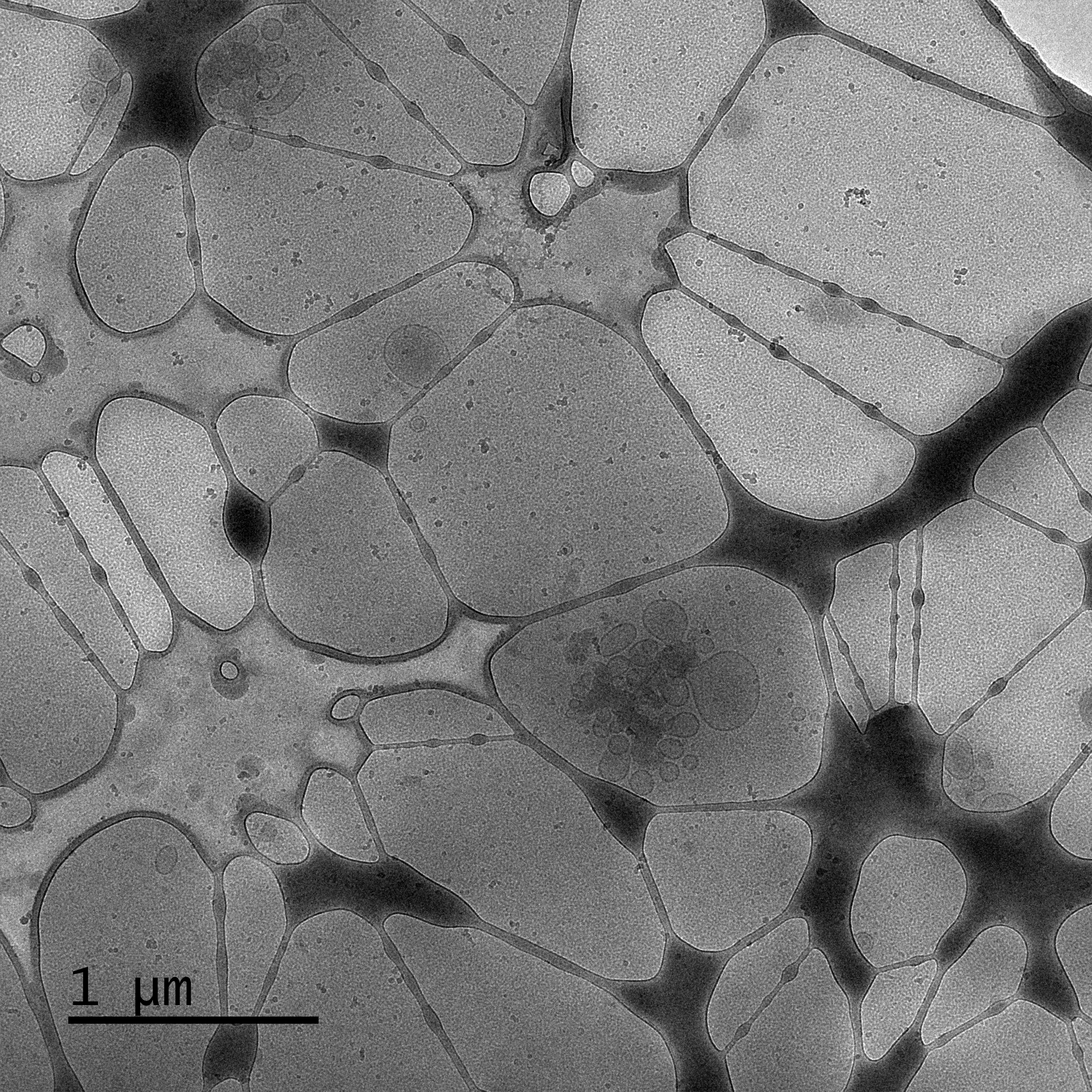

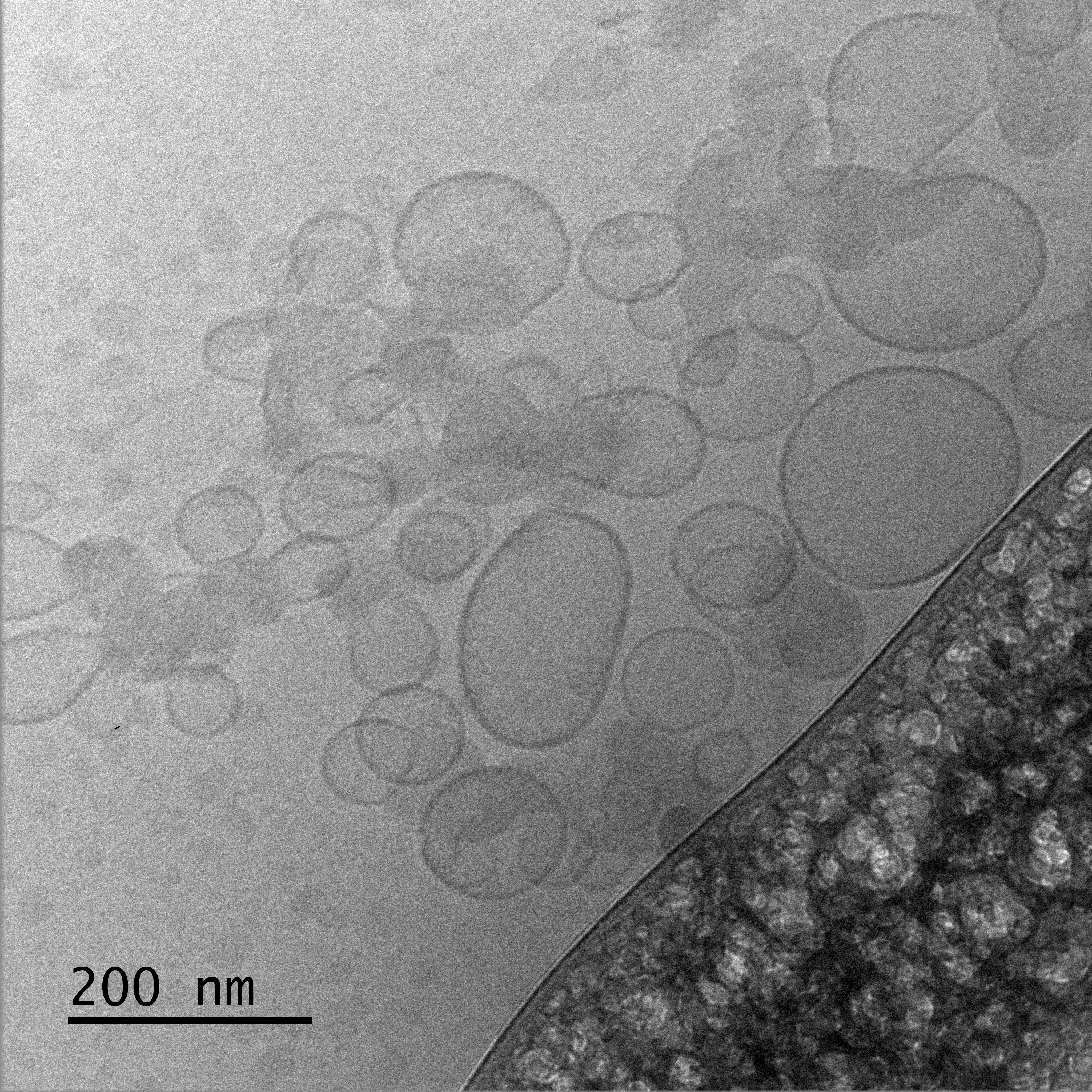

**Fig. S5. CyroEM of unmodified LNPs (upper panels) and DNA-LNPs (lower panels).** LNPs have a spherical morphology and maintain morphology after post-insertion of DNA-lipid.

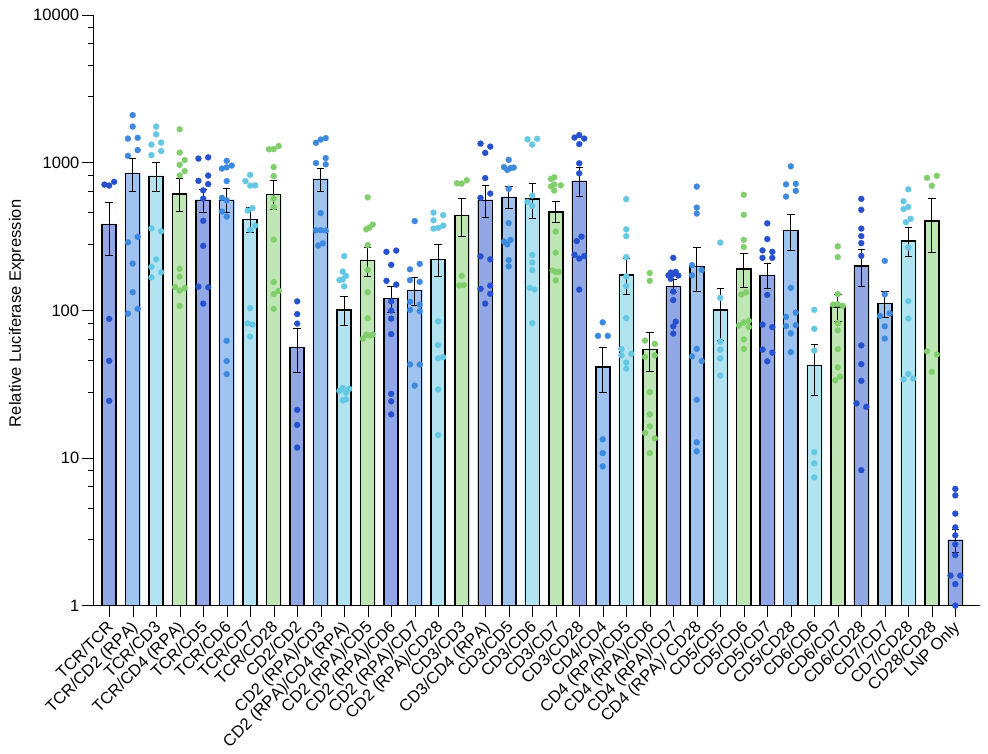

**Fig. S6. Jurkat cells expressing Fluc 24hrs after transfection.** Jurkat cells were transfected with 36 different tLNP formulations and efficiency of transfection is indicated by expression of firefly luciferase read as relative luminescence.

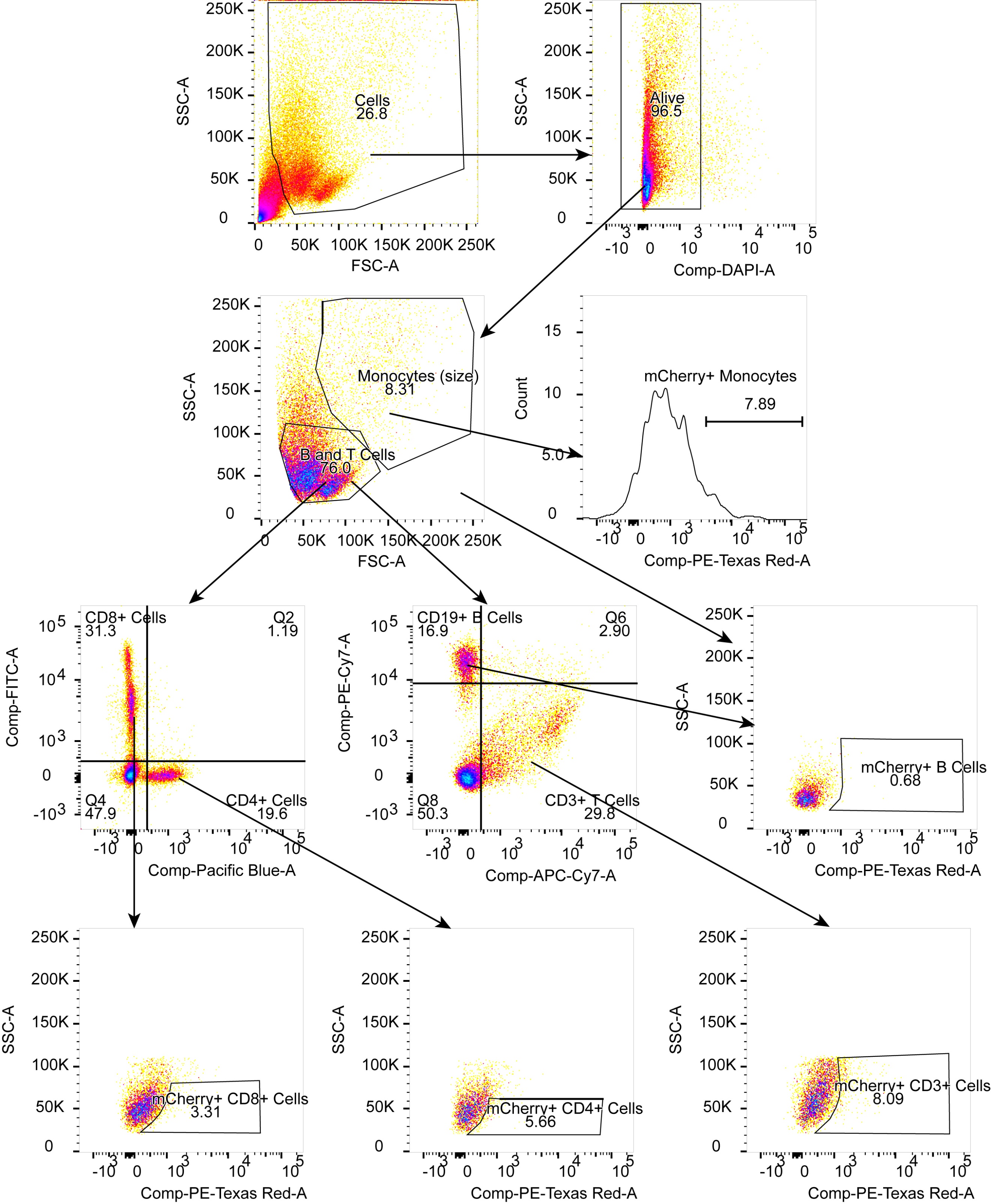

**Fig. S7. Flow cytometry gating strategy for human PBMCs.** Gating shown for PBMC with treated with CD3/TCR LNPs.

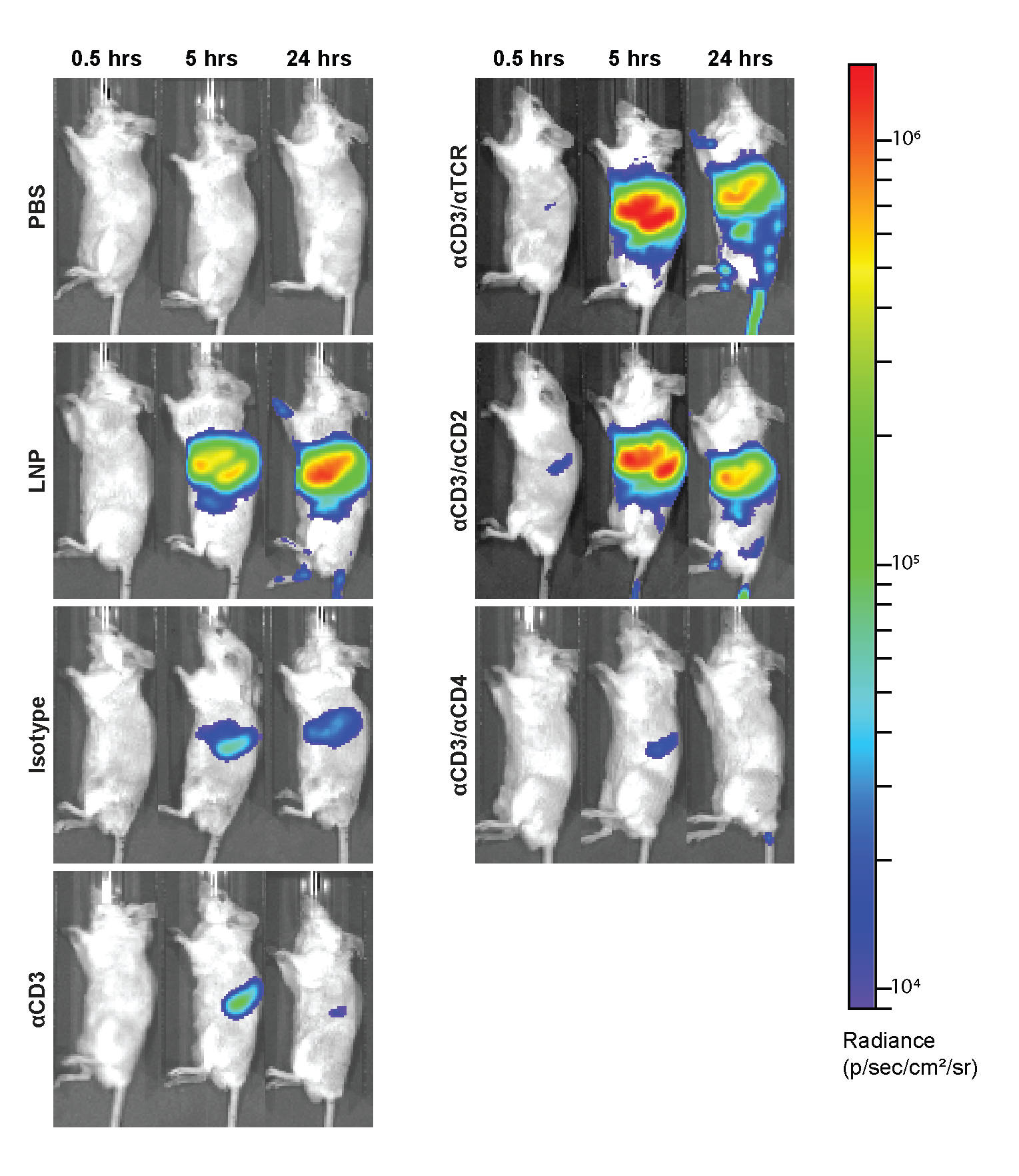

**Fig. S8. *In vivo* imaging of luminescence over time.** *In vivo* images of luminescence of the mice injected with Fluc-LNPs or controls were taken at time 0, 5, and 24 hours to examine distribution and intensity of transfection.

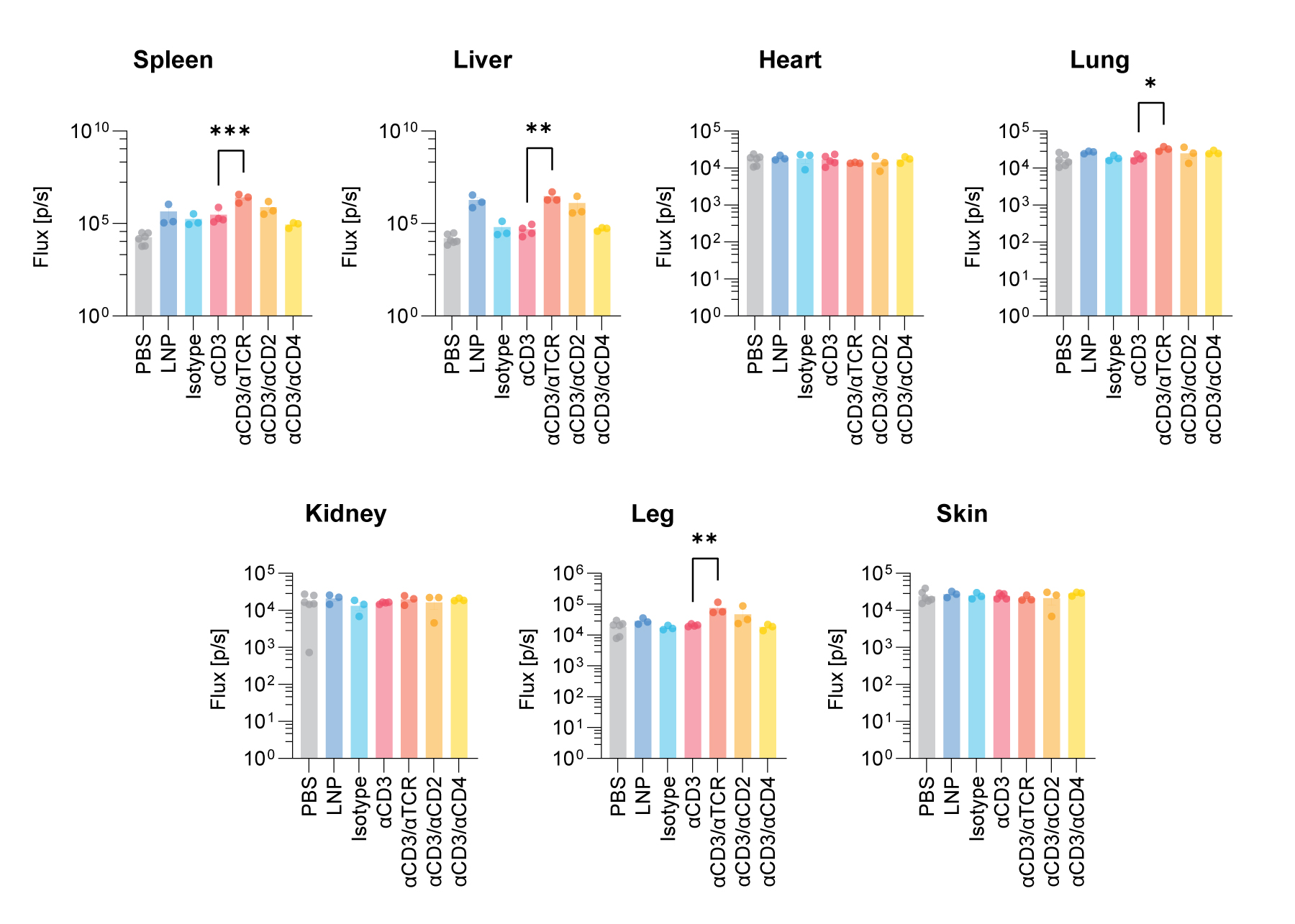

**Fig. S9. *Ex vivo* organ luminescence at 24 hours.** Organs were isolated 24 hours after injection with Fluc-LNPs or PBS control and imaged on the IVIS. Luminescence of organs is recorded as flux and indicated firefly luciferase expression.

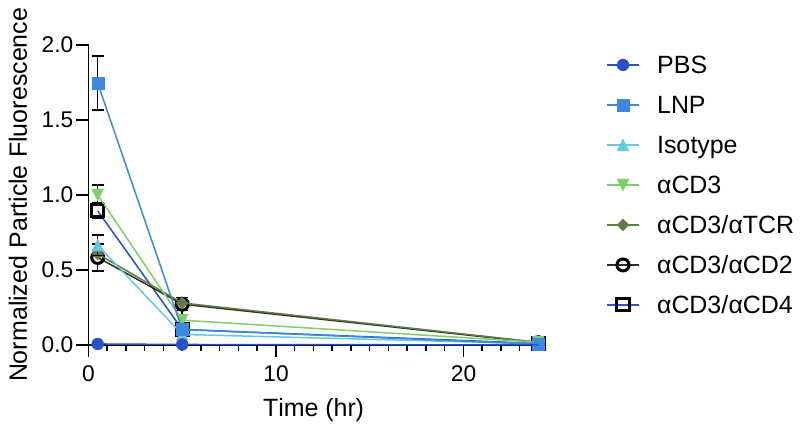

**Fig. S10. Particle clearance from the blood at t = 0, 5, and 24 hours.** Blood drawn retro-orbitally was analyzed on a plate reader for LNP fluorescence of Cy7 dye at t = 0, 5, and 24 hours. All particles were cleared from the blood by 24 hours.

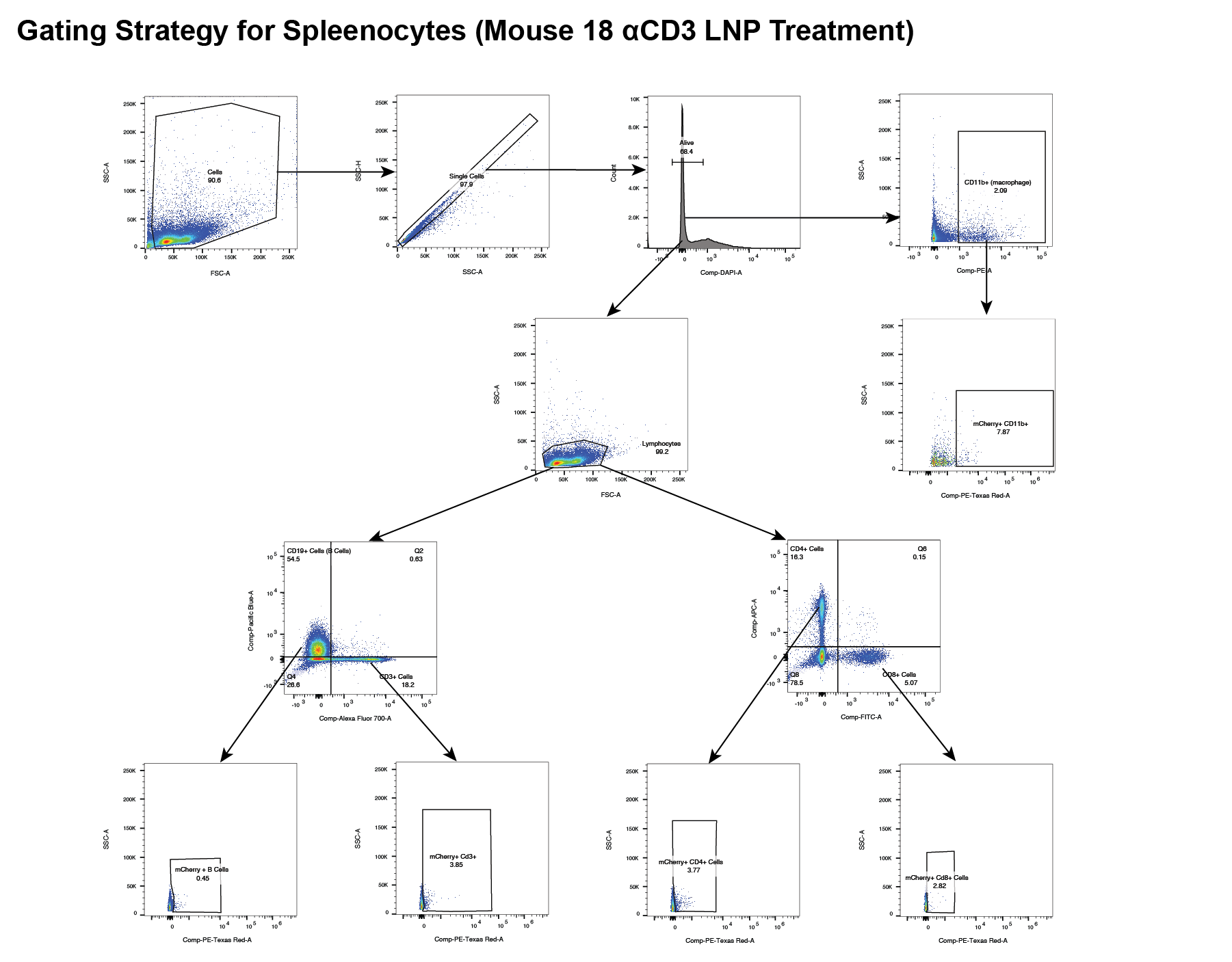

**Fig. S11. Particle clearance from the blood at t = 0, 5, and 24 hours.** Blood drawn retro-orbitally was analyzed on a plate reader for LNP fluorescence of Cy7 dye at t = 0, 5, and 24 hours. All particles were cleared from the blood by 24 hours.

Tables

Table S1. DNA sequences used for antibody tethering. DNA sequences with and without a spacer were tested for antibody attachment to LNPs.

|  | **Sequence** | **MW** |
| --- | --- | --- |
| **Z-DNA** | 5’- /5DBCON/GTA GTG CGC GAG GGT GA -3’ | 5,827 |
| **Z-teg DNA** | 5’-/5DBCOTEG/GTA GTG CGC GAG GGT GA -3’ | 5,902.1 |

Table S2. Nanoparticle tracking analysis of LNPs formulated with MC3 ionizable lipid. Mean particle size and mean concentration of LNPs carrying mCherry mRNA and Fluc mRNA cargos with no surface modification. Each value is the average of 2 biological replicates each with 5 technical replicates. SE = Standard Error.

|  | **Particle Size ± SE (nm)** | **Concentration ± SE (particles / mL)** |
| --- | --- | --- |
| **mCherry LNP** | 151.9 ± 1.8 | 4.06e^8^ ± 8.27e^6^ |
| **Fluc LNP** | 156.4 ± 3.4 | 2.27e^8^ ± 9.23e^6^ |
